# Fog adaptation and wind connectivity in an endangered California plant: a genomic baseline for wind energy development

**DOI:** 10.64898/2026.10.25.761743

**Authors:** Emma Steigerwald, Kristen Hasenstab-Lehman, Dorian Scheim, Milagros G Rivera, Matthew M. Kling, Zackary Werner, Susan McEvoy, C. Matt Guilliams, Rachel S Meyer

**Affiliations:** Department of Ecology & Evolutionary Biology, University of California, Santa Cruz; Santa Barbara Botanic Garden; Franke College of Forestry & Conservation, University of Montana; Independent Researcher; Department of Forest Genetics and Plant Physiology, Swedish University of Agricultural Sciences

## Abstract

Fog buffers water stress in several arid, highly seasonal coastal ecosystems with exceptional plant endemism, yet the genomic basis of adaptation to fog remains almost entirely uncharacterized. We resequenced 842 genomes from six localities of the endangered Gaviota tarplant (*Deinandra increscens* subsp. *villosa*), endemic to the fog-swept central California coast, where a wind farm was built across the largest population. Populations were strongly structured, and within the largest population, wind connectivity was weakly associated with genetic similarity, providing a pre-turbine baseline for gene-flow effects. Genotype–environment association analyses identified many fog-associated loci, and GO terms enriched among association signals clustered into functional themes including cell wall remodelling, osmoregulation, developmental timing, light perception, lipid and energy metabolism, and immunity. Independently, herbarium specimens show broader architecture and a higher proportion of ray florets in foggier, wetter sites, traits that fog-associated candidate genes may help explain. Among 31 fog-associated peaks, a centromere-associated haplotype showed the strongest response to germination-year fog. That response, together with elevated long runs of homozygosity and a smoother allele-frequency surface in 2021, an extreme drought year with unusually rich fog, is consistent with early-season environmental filtering. This baseline informs management and motivates germination and survival trials to test early-season filtering.

## Introduction

Along the California coast, persistent summer fog and low clouds moderate the seasonal extremes of the Mediterranean climate, creating microclimates with distinct water availability and temperature. Fog can benefit plants through fog drip and foliar water uptake, reduced evaporative loss, nutrient deposition, and sustained photosynthesis (Cuevas *et al*. 2023; Glaser *et al*. 2022; Gomes & Baguskas 2022; Lehnert *et al*. 2018; Sun *et al*. 2024), but it can also impose costs, including reduced light and enhanced microbial activity (Evans *et al*. 2018; Mieslerová *et al*. 2022; Ritter *et al*. 2009). This balance of benefits and costs may generate spatially variable selection along fog gradients.

Gaviota tarplant (*Deinandra increscens* subsp. *villosa*; Madiinae, Asteraceae) offers an opportunity to study local adaptation to variable fog in a range-restricted California endemic. This federally endangered annual herb occurs in a few populations spanning Point Conception (USFWS 2022). One of the few plants flowering between June and November, it is a critical resource for pollinators (Cusser et al. 2024) and was traditionally harvested by the Chumash, who are thought to have burned grasslands to enhance tarweeds and other valued plants (Timbrook *et al*. 1982). Summer rainfall is largely absent where it grows, but many coastal California plants offset this deficit with fog drip or foliar uptake (Kennedy & Sousa 2006; Fischer *et al*. 2009), and some coastal grassland species obtain two-thirds of their summer water from fog (Corbin *et al*. 2005). The tarplant’s habitat and morphology are consistent with fog specialization: it occupies fog-exposed coastal bluffs, flats, and topographic saddles in grassland and scrub, and has dense pubescence, narrow leaves, abundant glandular trichomes, and a highly branched architecture (Konrad *et al*. 2015; Martorell & Ezcurra 2007; Stanton & Horn 2013; USFWS 2022). Fog dependence is well documented in woody and other perennial plants, but its significance for annual plants remains poorly characterized (Weathers *et al*. 2020).

Remaining Gaviota tarplant populations face rapid landscape conversion from agricultural, urban, residential, and military development (Shapero *et al*. 2022) and pronounced declines in coastal fog over the past century (Johnstone & Dawson 2010). The 24 km² Strauss Wind Farm was recently built across the subspecies’ largest remaining population (USFWS 2022). Turbines reduce near-surface wind speed, decrease humidity, accelerate evaporation, and may alter fog (Akhtar *et al*. 2024; Adkins & Sescu 2022; Hasager *et al*. 2023; Liu *et al*. 2022), which could affect gene flow if wind influences the movement of the plant’s bee and beetle pollinators (Cusser *et al*. 2024) and cause adaptive mismatch if turbines shift the fog conditions to which plants may be locally adapted. A genomic and phenotypic baseline is therefore an urgent priority, and major collaborative conservation efforts are underway to inform management.

Here, building on a recently published reference genome notable for its complex evolutionary history of repeat activity (McEvoy *et al*. 2024), we use whole-genome resequencing of 842 individuals across six populations to characterize genomic variation across the range of Gaviota tarplant and investigate its relationships with environmental heterogeneity. We first characterize patterns of genetic diversity and population structure among populations spanning the subspecies’ geographic range and assess wind connectivity as a potential driver of gene flow, establishing a baseline against which future wind farm-induced changes can be evaluated. We then test for genomic and phenotypic signatures of local adaptation, with particular emphasis on fog, using association analyses and a century-long record of herbarium specimens. We aim to provide both insight into the evolutionary processes shaping environmental specialization and a framework for incorporating genomic information into conservation monitoring and management.

## Results

### Strong population structure and geographic variation in genetic diversity

Gaviota tarplant populations are strongly structured, with patterns of differentiation that are not well explained by geographic distance (five populations: Mantel *r* = 0.45, *p* = 0.27; including Jalama: *r* = 0.25–0.39, *p* = 0.28–0.31). PCA and admixture analyses both reveal clear compositional differences among localities, with structure continuing to emerge up to *K* = 7 (**Fig. 1B,C**; **Fig. S1**). Most pairs of sampling localities show a moderate degree of genetic differentiation (*F_ST_*= 0.05–0.08; Fig. 1D). However, Gaviota is highly differentiated from every other locality (*F_ST_* = 0.13–0.15), a pattern that persisted when all populations were downsampled and coverage-matched. Jalama also appeared highly differentiated in full-sample estimates (*F_ST_* = 0.09–0.18), but this reflected the low sequencing coverage of most of its samples: its excess differentiation disappeared when the other populations were coverage-matched to it.

**Figure 1.**
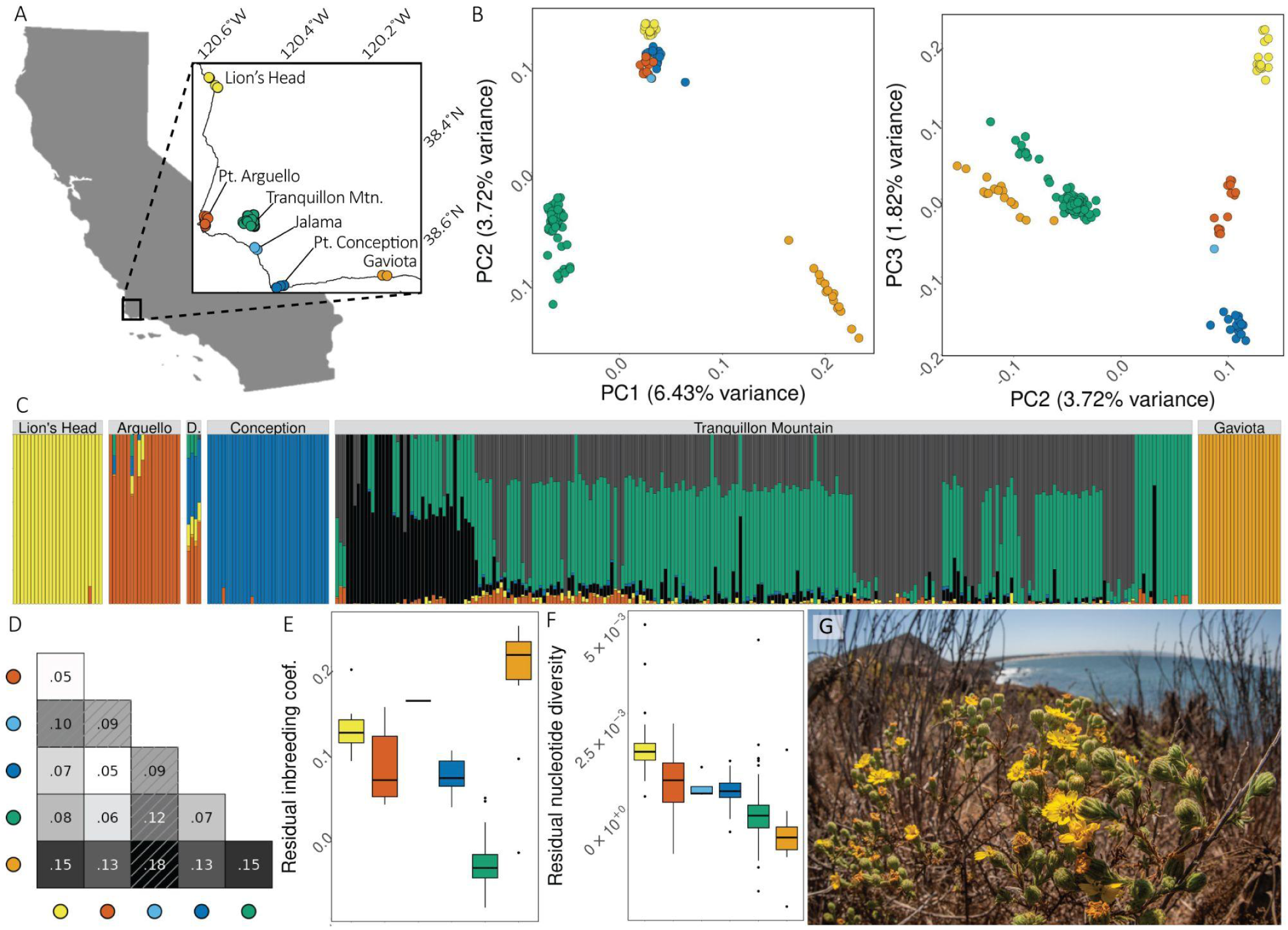
Genetic structure and diversity by localities. (A) Sampling map of Gaviota tarplant populations as an inset, showing the wider context for the study area in California. (B) Principal component analysis of genomic variance, with individuals colored by the locality from which they are sampled. (C) OHANA analysis of ancestral components at *K* = 7 components. Facets sort samples by locality, and within facets samples are sorted by longitude. The predominant ancestral component within a locality is colored to match the color assigned in panel A. (D) Pairwise *F_ST_* between populations; Jalama values (hashed) are inflated by low sequencing coverage. (E) Individual inbreeding coefficients and (F) nucleotide diversity, corrected by coverage, colored by locality as in panel A. (G) Gaviota tarplant at the Lion’s Head locality atop coastal bluffs. Photo by E. Steigerwald.

Inbreeding coefficients differ significantly among populations (*Kruskal–Wallis χ²* = 115.16, *df* = 5, *p* < 0.001), with the more differentiated Jalama and Gaviota populations also showing the highest inbreeding (**Fig. 1E**). Tranquillon Mountain has significantly lower inbreeding than Gaviota, Point Conception, Point Arguello, and Lion’s Head (*adjusted p* < 0.001), consistent with its high census size (**Table 2**). Calculated at the population scale, nucleotide diversity declines from 7.2 × 10^-3^ for Lion’s Head at the northern extent to 5.8 × 10^-3^ for Gaviota to the southeast (**Table 2**), placing *Deinandra increscens* subsp. *villosa* well below the exceptionally high values reported for other obligately outcrossing species (**Fig. S2**; Leffler *et al*. 2012). Individual expected heterozygosity corrected for autosomal coverage varies significantly among populations (*Kruskal-Wallis χ²* = 147.83, *df* = 5, *p* < 0.001; **Fig. 1F**).

**Table 2.** Nucleotide diversities (π), effective population sizes (N_e_), and census population sizes of Gaviota tarplant populations. Census population sizes are taken from USFWS (2022).

| **Locality** | **π** | N<sub>e</sub> | **Census population size** |
|---|---|---|---|
| Lion's Head | 7.2 × 10<sup>-3</sup> | 460,000 | 611 |
| Pt. Arguello | 6.7 × 10<sup>-3</sup> | 408,200 | 750 |
| Jalama | 6.2 × 10<sup>-3</sup> | 346,800 | 58,000 |
| Pt. Conception | 6.6 × 10<sup>-3</sup> | 355,000 |  |
| Tranquillon | 6.3 × 10<sup>-3</sup> | 334,100 | 6,664,767 |
| Gaviota | 5.8 × 10<sup>-3</sup> | 227,800 | 12,000 |

### Fine-scale structure and a drought-year signature in the largest population

Within Tranquillon Mountain there was fine-scale genetic structure (**Fig. 2A-B**, **Fig. S3**) and a real but weak signal of continuous isolation by distance (*Mantel r* = 0.119, *p* < 0.0001; **Fig. S4**). There is also support for both spatial and temporal patterns in inbreeding within the Tranquillon Mountain locality. The highly-structured, western portion of Tranquillon Mountain is characterized by higher inbreeding coefficients than the eastern portion (**Fig. 2C**). Proportion of the genome in long (>1 Mb) runs of homozygosity differed significantly among years (Kruskal-Wallis *χ²* = 9.92, *df* = 2, *p* = 0.0070), and long ROH was significantly elevated during the extreme drought year (2021) compared to non-drought years (2020 and 2022 combined; *Wilcoxon rank sum test*, *W* = 18270, *p* = 0.0022; **Fig. 2D-E**).

**Figure 2.**
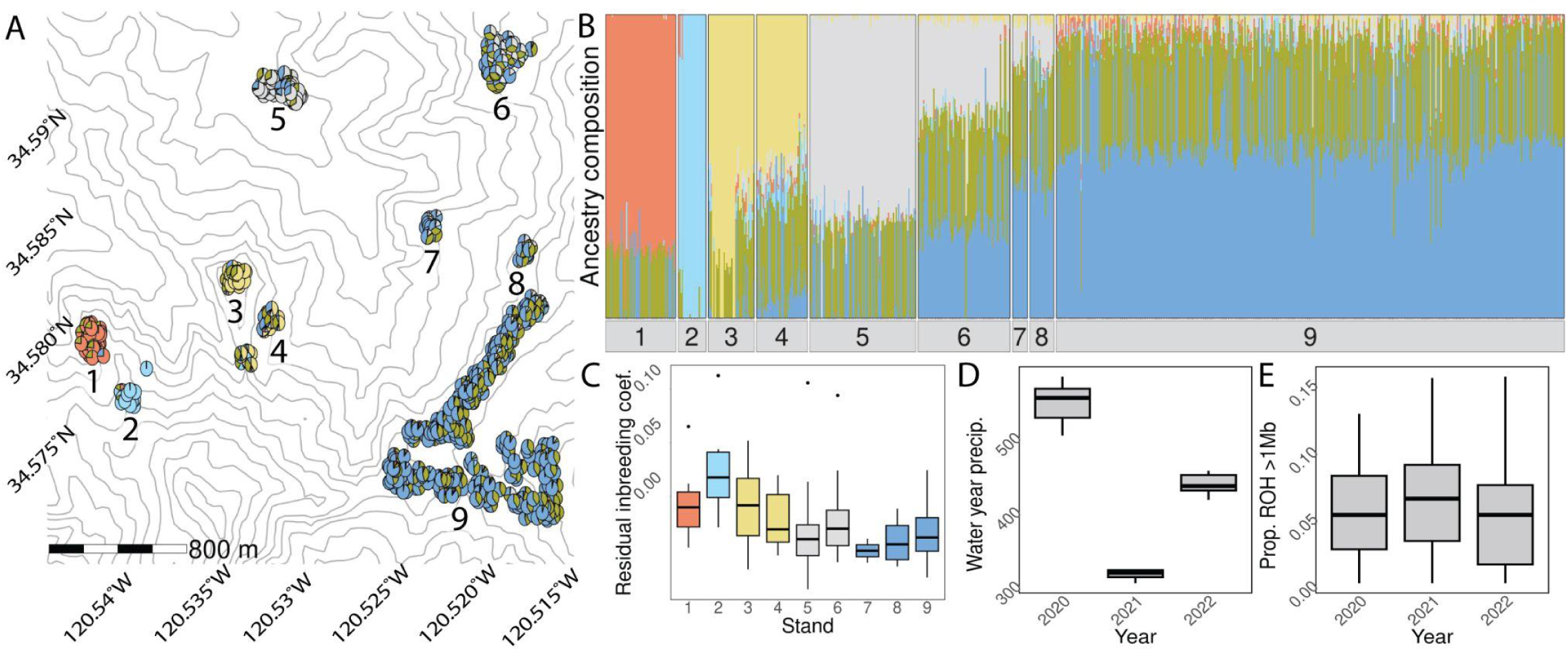
Genetic structure within the Tranquillon Mountain locality. (A) OHANA analysis of ancestral components at *K*=6 components, projected as pie charts onto a contour map of the Tranquillon Mountain site. Stands of plants are labelled 1-9. Wind turbine placement is indicated with black icons. (B) OHANA ancestry components shown split into panels by stands. (C) Inbreeding coefficients for stands of plants in Tranquillon Mountain, colored to match the dominant admixture component as shown in panels A/B. (D) Proportion of the genome in long (>1 Mb) runs of homozygosity (ROH) for Tranquillon Mountain plants across three years of sampling, where 2021 represented an extreme drought year.

The western portion of Tranquillon Mountain was characterized by lower migration surface weights, consistent with the higher inbreeding and stronger genetic structure observed in that portion of the locality (**Fig. S5**). The optimal lambda differed among cohorts: during an extreme drought year in 2021 (**Fig. 2D**), we see a lambda (*λ* = 452.3) approximately 1.79-fold higher than the 2020 cohort (253.1) and 4.70-fold higher than the 2022 cohort (96.2). This higher lambda reflects a more homogeneous allelic composition across Tranquillon Mountain in the extreme drought year.

### Wind connectivity is weakly associated with fine-scale genetic differentiation

To ask whether wind structured range-wide differentiation, we retained only five widely separated populations (Jalama’s F_ST_ being inflated by low coverage), so power was low. Pairwise F_ST_ was unrelated to wind connectivity after conditioning on geographic distance, environmental distance, or both (*all p* ≥ 0.39; **Fig. S6A,B**), including when connectivity was modeled as a random walk with airborne half-lives from 0.5 to 24 h (*all p* ≥ 0.26), and when Jalama was included (*all p* ≥ 0.16). Because prevailing winds blow from the northwest, wind-mediated gene flow would be expected to make downwind (southern) populations more genetically diverse than upwind ones (Kling & Ackerly 2021). Given the north-to-south decline nucleotide diversity reflecting the decline in effective population sizes (**Table 2**), this test cannot separate wind from other north-to-south gradients, and populations receiving more wind flow from a partner than they sent to it tended instead to be less diverse than that partner (*r* = −0.70, p = 0.11; random-walk connectivity: *r* = −0.82 to −0.83, *p* = 0.067; with Jalama: *r* = −0.60 to −0.82, *p* = 0.053–0.15).

Within Tranquillon Mountain, a site under 4 km across, where net wind flow varied from 1.6 to 3.6 m s⁻¹ and its direction spanned 44° across sampled plants, we found weak evidence that wind connectivity shapes genetic differentiation. Individuals linked by weaker wind flow were slightly more differentiated than expected from distance alone (*r* = 0.045, *p* = 0.047; *n* = 276; **Fig. S6C,D**) or from distance and environment together (*r* = 0.045, *p* = 0.042). The association was of similar strength in every WindNinja configuration, from terrain-only wind (*r* = 0.049, *p* = 0.037) to versions adding a finer mesh, spatially varying vegetation, diurnal slope flows or non-neutral atmospheric stability separately (*r* = 0.047–0.050, *p* = 0.034–0.041). Gene flow was not biased in the downwind direction: the asymmetry of gene flow along grid edges near sampled plants was unrelated to the asymmetry of wind flow (*r* = 0.16, *p* = 0.32; *r* = 0.10–0.18, *p* = 0.13–0.60 across smoothing values).

### Genomic signatures of local adaptation to fog regime

Despite weak-to-moderate intercorrelations among fog, precipitation seasonality, and summer precipitation (*maximum pairwise Spearman’*s *r* = 0.54), their associated outlier peaks and candidate gene sets showed substantial overlap (**Figure 3A**; **Fig. S7**). Of the 161 candidate genes identified across these variables (**Table S1**), 39% were shared by at least two variables, and 7% were shared by all three. (hypergeometric tests, all *p* < 10⁻²⁵; **Fig. S8**). Enriched GO terms also converged modestly but non-randomly between environmental variables (**Fig. S9**).

**Figure 3.**
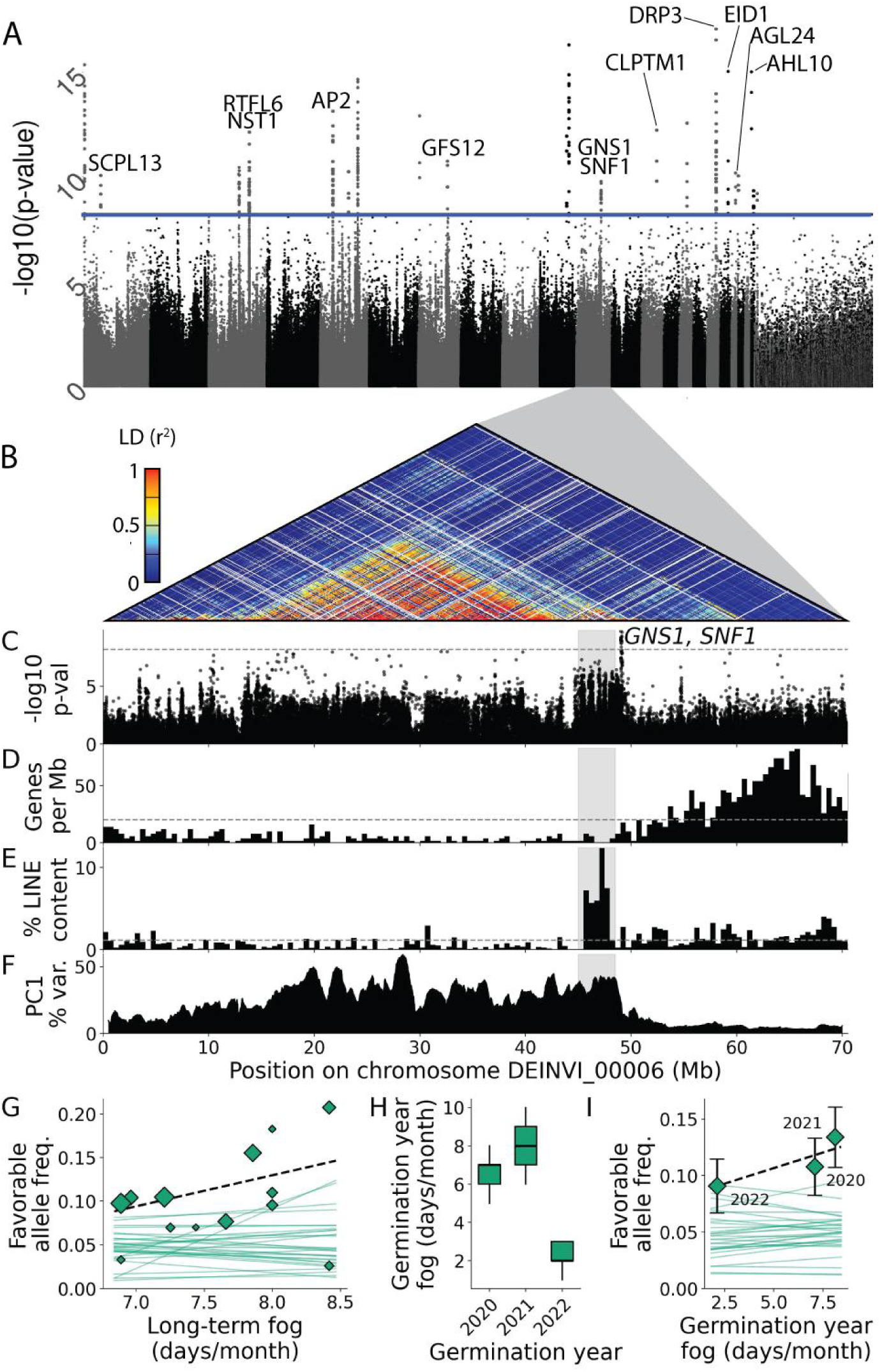
Local adaptation to fog exposure. (A) GxE results for fog. Scaffolds (alternating grey/black) shown in descending size order. Peaks are labelled with candidate genes; unlabelled peaks have no annotated gene within 5 kb of, or in linkage disequilibrium (*r²* ≥ 0.5) with, their outlier SNPs. The blue line is the Bonferroni threshold. (B) Linkage disequilibrium across chromosome DEINVI_00006. (C) GxE results for fog for DEINVI_00006. The dashed line is the Bonferroni threshold. (D) Gene density and (E) percent LINE content in 500-kb windows. Dashed lines give genome-wide average values. (F) Percentage of genetic variance explained by PC1 in sliding-window PCA. (G) Favorable-allele frequency for stands of Tranquillon Mountain plants against the long-term fog exposure, where the favorable allele at each SNP is the one whose coverage-adjusted dosage rises with fog. Diamonds show values for the DEINVI_00006 peak, with area proportional to the number of plants in the stand, and the dashed line representing the *n*-weighted least-squares fit. Thin lines show the same fits for the other 30 fog outlier peaks. (H) June fog in each plant’s germination year. (I) Favorable-allele frequency per germination cohort, adjusted for stand, against that cohort’s mean June fog. Diamonds show the DEINVI_00006 peak, with error bars showing the 95% CIs and the dashed line fitted as the within-stand slope. Thin lines show the same for the other 30 fog-outlier peaks.

Association analysis identified 443 variants significantly associated with fog exposure, organized into 31 genomic peaks distributed across 12 scaffolds. The enriched GO terms associated with fog exposure (FLCC; raw *p* < 0.005; none FDR-significant) largely fell into a small number of coherent functional groups consistent with the morphological and physiological demands of a fog-exposed coastal environment (**Fig. S9A**): cell wall and carbohydrate remodeling (GO:0016759, GO:0000272, GO:0031501, GO:0015925), altered light perception (GO:0090227, GO:0009584), osmoregulation and leaf surface traits (GO:0010325, GO:1904278), immune function (GO:0051502), developmental timing (GO:0010331, GO:0045962), and lipid and energy metabolism (GO:0010896, GO:0004806, GO:0072350). Although no GO term reached FDR significance, a permutation test rules out test-statistic artifact as an explanation: the observed analysis yielded 18 genome-wide-significant SNPs for fog exposure, compared with a permuted-null mean of 0.94 (*range* 0–17; *empirical p* = 0.0099). All fog-nominated candidate genes showed detectable expression (TPM > 0) in at least one tissue type, confirming that the annotated loci are actively transcribed in *D. increscens* subsp. *villosa* (**Fig. S10**).

### A centromere-associated haplotype tracks fog in space and time

Among the fog-associated outlier peaks, one on scaffold DEINVI_00006 is distinguished by its centromeric genomic context. The peak of 19 SNPs contains two genes and lies at the edge of a broad region of elevated association and high LD spanning 44–49 Mb, which is itself adjacent to the largest recombination-suppressed haploblock in the genome spanning 16-44Mb (**Fig. 3B,C; Table S1**). This association did not reflect a mapping artifact (Methods). LINE content rises sharply in a single, well-bounded interval at 45.5–48.0 Mb (**Fig. 3E**), and forms a tight cluster on nearly every chromosome-scale scaffold, mirroring the centromere-marking LINE-like repeats of sunflower (**Fig. S11**; Nagaki *et al*. 2015). This interval is gene-poor, with density rising toward the chromosome arms (**Fig. 3D**). Population-genetic signatures are consistent with a non-recombining centromeric region and argue against a chromosomal inversion: windowed PCA shows elevated PC1 variance from ∼16–49.3 Mb with individuals falling into three discrete groups (**Fig. 3F**; **Fig. S12B,C**), and LD within each group is 3–7 times higher than in control regions of the same chromosome, whereas an inversion would suppress recombination only in heterozygotes. While the outlier peak lies only 0.59 Mb from the edge of the centromere, fog outlier peaks as a whole show no bias toward centromeres (*p* = 0.29), nor do outlier peaks for other climate variables (*p* ≥ 0.47; **Fig. S11**).

Comparative synteny places the DEINVI_00006 centromere at an ancestral chromosomal junction. Alignment with sunflower (*Helianthus annuus*) and lettuce (*Lactuca sativa*) places the left arm and right arm in entirely different ancestral chromosomes, with no syntenic genes spanning the centromeric interval (**Fig. S12D**). The two candidate genes at the fog outlier peak, GNS1 and SNF1, are 22 kb apart in *D. increscens* but 1.0–1.7 Mb apart in sunflower & lettuce, with ∼23 intervening genes. Those intervening genes retained in D. increscens lie on a homologous region of DEINVI_00020, which, with DEINVI_00006 and 00011, maps in collinear, interleaved order along the lettuce segment—a signature of differential gene loss following the genome triplication shared between sunflowers and *Deinandra* (McEvoy et al. 2024) rather than a recent rearrangement. Together, these findings indicate that the unusual population-genetic properties of this haplotype reflect recombination suppression rooted in chromosome evolution.

The DEINVI_00006 haplotype tracks fog at both spatial and temporal scales within Tranquillon Mountain, ranking 3rd and 1st of 31 fog-outlier peaks respectively. Across the subspecies’ range, the favourable haplotype is common only at the foggiest site (Point Arguello, 0.66) and rare or absent elsewhere (0.00–0.24), so the landscape-scale association reflects largely a single locality contrast; how it tracks fog within Tranquillon Mountain was therefore tested. For the spatial test, DEINVI_00006 haplotype frequency rose 0.036 per fog day/month across stands (**Fig. 3G**), though the gradient was not significant against permutation of fog among stands (*p* = 0.080), frequency-matched SNPs (*p* = 0.106), or centromere-distance-matched SNPs (*p* = 0.098). This test is not independent of peak discovery — favourable alleles were defined by long-term fog in a sample 82% composed of Tranquillon Mountain plants, yet 15 of the other 30 peaks showed zero or negative slopes, suggesting most fog associations reflect between-locality contrasts. For the temporal test, the DEINVI_00006 haplotype rose 0.0055 per fog day/month with germination-year fog within stands (**Fig. 3I**; **Fig. S13**). This was significant against permutation of germination-year fog within stands (*p* = 0.025) and frequency-matched SNPs (*p* = 0.029), but marginal against centromere-distance-matched SNPs (*p* = 0.054). Unlike the spatial test, this comparison is independent of peak discovery, which uses long-term rather than year-to-year fog.

The fog outlier peak lies 0.59 Mb from the DEINVI_00006 centromere— a classic site of female meiotic drive— raising the question of whether drive rather than selection explains its signals. Against random blocks at the same distance from their own centromeres, the peak’s Tajima’s D was unremarkable in four of five populations (31st–73rd percentile) and low only at Pt. Arguello (5th percentile), the one locality where the fog haplotype is common (0.66; **Fig. S14**). Centromeric haplotype frequencies, differentiation, and heterozygote deficit were inconsistent with a sweep-like driver (**Supplementary Note**). These checks do not exclude balanced drive, in which a costly driver persists at intermediate, locally variable frequencies (as for the *Mimulus guttatus* D locus; Fishman & Saunders 2008), which would require transmission data to test.

### Tarplant architecture and floral investment respond to fog & summer precipitation

Gaviota tarplant morphology varies significantly among localities and responds to the same environmental gradients implicated in genomic adaptation. In the modern dataset, greater summer moisture is associated with a lower height:width ratio (fog: *β*= −0.67 ± 0.176, *t* = −3.80, *p* < 0.001; summer precipitation: *β* = −2.04 ± 0.885, *t* = −2.30, *p* = 0.022; *n*= 115; **Fig. 4B**; **Table S2**) and a higher ray to total floret ratio (fog: *β* = 0.052 ± 0.025, *z* = 2.10, *p* = 0.036; summer precipitation: *β* = 0.067±0.033, *z* = 2.05, *p* = 0.041; *n* = 138; **Fig. 4C**; **Table S2**).

**Figure 4.**
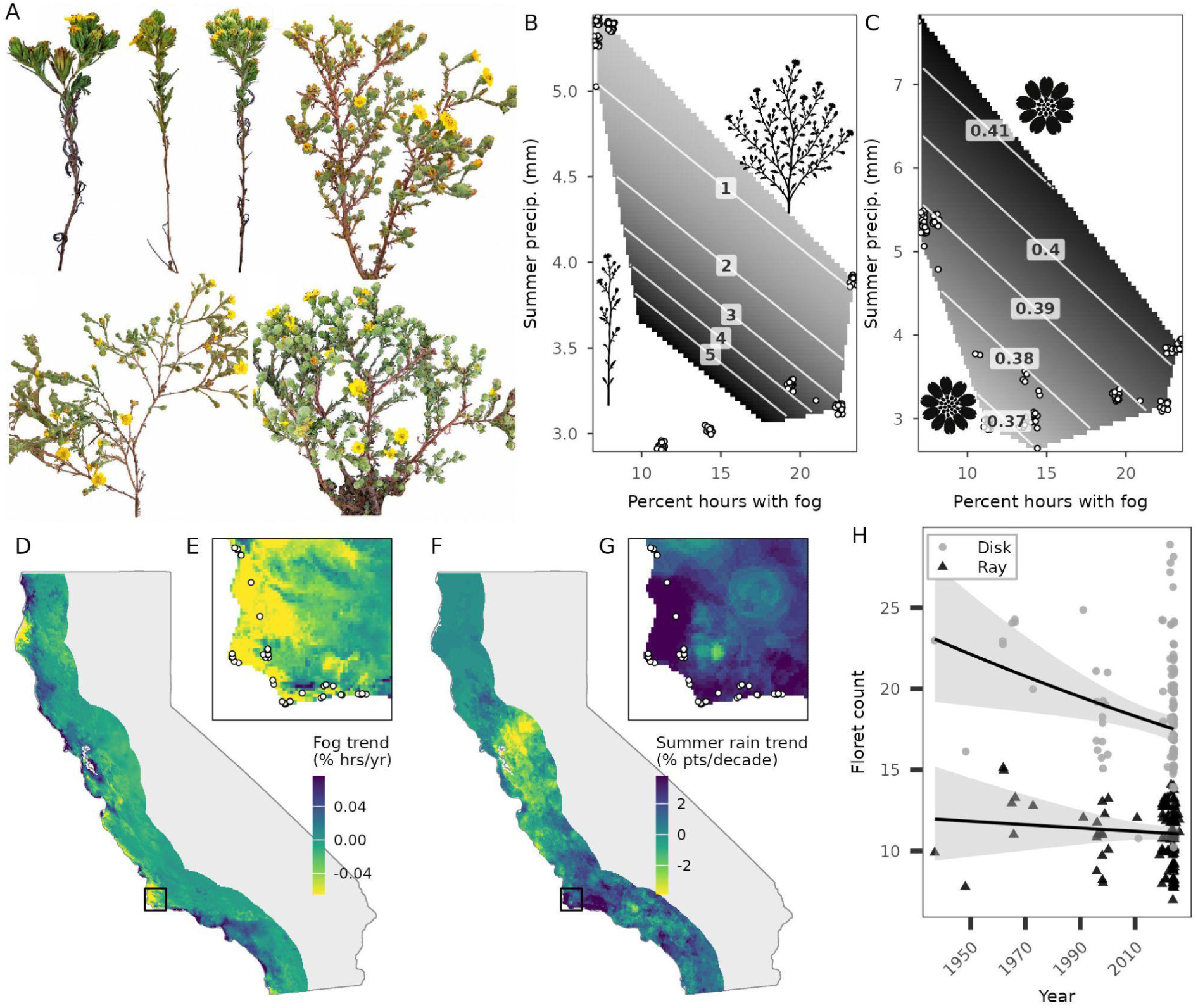
Phenotypic and environmental change in the Gaviota tarplant across the modern and historic sampling periods. (A) Plants representing extreme high (first three) and low (last three) values of height:width ratio. (B-C) Response surfaces showing how plant height:width ratio (B) and proportion of ray florets (C) vary jointly with fog frequency and summer precipitation; points are observed specimens (jittered), contour lines show model-predicted values, and silhouettes illustrate the phenotypic extremes at each surface’s corners. Surfaces are clipped to the observed range of each trait. (D-E) Trends in the percentage of hours with fog from 1950-2014 for (D) the California coast and (E) the study region. (F–G) Trends in the probability of measurable summer rain, 1927–2025, for (F) the California coast and (G) the study region, expressed as the change in percentage points per decade. (H) Disk (grey circles) and ray (black triangles) floret counts by collection year, historic to modern record, with population-level GLMM trend lines and 95% CI.

The environmental gradients associated with Gaviota tarplant’s floral morphology have shifted substantially over the historic record: fog frequency declined significantly (*Mann-Kendall τ* = −0.21, *p* = 0.013; **Fig. 4D–E**) and summer rain occurrence showed a marginally significant increase (*τ* = 0.14, *p* = 0.056; **Fig. 4F–G**). Disk floret count has declined significantly since 1927 (*β* = −0.049 ± 0.018, *t* = −2.73, *p* = 0.006, *n* = 162; observed range 8–29), with no clear change in ray floret count (*n* = 213; observed range 7–15; **Fig. 4H**) and a consequent marginal increase in ray proportion (*β* = 0.034 ± 0.019, *z* = 1.8, *p* = 0.072, *n* = 162). The direction of this morphological shift is consistent with the increase in summer precipitation, but not with the decline in fog.

## Discussion

Coastal fog is a defining feature of endemic-rich terrestrial ecosystems worldwide—from California’s floristic province to Namib Desert fog-basking communities and montane cloud forests—yet fog regimes in many of these systems are declining under climate change (Foster 2010, Lavigne & Liu 2022, Li *et al*. 2025). The genomic basis of plant adaptation to fog has, to our knowledge, not previously been characterized. Here we characterize adaptation in the Gaviota tarplant by bringing together genomic and phenotypic data, climate models, remotely-sensed and machine-learning fog products, and in situ plant data. This advances understanding of how plants adapt genomically to fog gradients and provides a conservation baseline for monitoring this subspecies as its environment changes and for identifying genotypes best positioned to persist.

### A weak role for wind in gene flow, with implications for wind energy development

The Strauss Wind Farm, which began operating in 2023 across the largest remaining population of Gaviota tarplant, changes the wind environment of this subspecies, and our 2020–2022 samples and pre-turbine wind climatology let us ask whether wind shaped gene flow beforehand. Range-wide, wind connectivity was unrelated to differentiation, but with only five widely separated populations, each shaped by drift and local adaptation, power was low and the null result is weakly informative. Within Tranquillon Mountain, a site under 4 km across, we expected at most a weak effect and found one: plants linked by weaker wind flow were slightly more differentiated than expected from distance and environment (*r* ≈ 0.05), with no detectable downwind bias.

Gaviota tarplant pollen is thought to be moved mainly by insects: its most frequent visitors are soft-winged flower beetles (Dasytinae; 34% of visits), sweat bees (*Lasioglossum* (*Dialictus*) spp.), blister beetles (*Epicauta* sp.), and long-horned bees (*Melissodes* sp.; Cusser *et al*. 2024). These species likely move only hundreds of metres, judging from studies of solitary bees and related blister beetles (Gathmann & Tscharntke 2002; Kendall *et al*. 2022; Marschalek & Young 2013; Marschalek *et al*. 2020); movement distances have not been reported for Dasytinae (Mawdsley 2003). Insect flight is also influenced by wind but not confined to downwind travel (Cardé *et al*. 2021; Combes *et al*. 2023). Seeds likely move by two routes: disk achenes bear a short, scaly pappus that could result in limited wind dispersal (USFWS 2022), whereas ray achenes lack one and are probably carried by animals with their involucral bract (Tanowitz *et al*. 1987). Wind-borne seed movement is therefore probably short-range, and our analyses cannot separate it from pollen movement. Because wind turbines reduce near-surface wind speed and add turbulence (Baidya Roy *et al*. 2004; Abraham & Hong 2021), and pollinator–flower interactions change near operating turbines (Weschler & Tronstad 2024; Weschler *et al*. 2026), the Strauss Wind Farm could alter gene flow by changing pollinator or seed movement. Repeating these analyses with post-construction samples would test this directly.

### Drought-year filtering may enrich fog-adaptive genotypes

The co-occurrence in 2021 of elevated long ROH, a smoother migration surface, and enrichment of the fog-adaptive centromeric haplotype is consistent with environmental filtering during an extreme drought year where fog represented an unusually important source of moisture. Two mechanisms could produce this pattern: early-season mortality of drought-sensitive seedlings, or differential germination by genotype. Distinguishing filtering from reduced effective dispersal under drought or stochastic cohort sampling would require demographic modeling or experimental germination trials across genotype classes. If filtering did occur, seeds collected during extreme drought years may be enriched for stress-tolerant genotypes better able to exploit fog as an alternative water source, a consideration for seed banking. However, it is important to note that elevated inbreeding may itself reduce drought fitness (Hauser & Loeschcke 1996; Lloret & García 2016), suggesting the picture is unlikely to be simple.

### Fog adaptation is polygenic, implicating the plant’s physical water interface and its environmental sensing machinery

Fog-associated candidate loci point to six nonexclusive functional themes that suggest hypotheses for how fog-related environmental variation could shape plant traits. Several fog-associated functional themes concern the plant’s physical interface with water and wet environments. Candidate loci implicate cell wall and carbohydrate remodeling, with NST1 as a notable candidate: in *Arabidopsis* and cotton, NST1 is a key regulator of secondary-wall formation and thickening (Mitsuda *et al*. 2007; Zhang *et al*. 2025). Osmoregulation and leaf surface traits also feature prominently, consistent with the importance of cuticle composition for fog water interception, transpirational water loss, and cellular stress tolerance under prolonged leaf wetness (Chin *et al*. 2022); candidate gene ANL2, for example, shows reduced leaf cutin when mutated in *Arabidopsis* (Nadakuduti *et al*. 2012). A third theme concerns immune function, which may reflect elevated microbial pressure in persistently wet environments (Evans *et al*. 2018; Mieslerová *et al*. 2022): one fog-associated candidate encodes a CYP76-family cytochrome P450, and in rice, related CYP76M enzymes including CYP76M7 contribute to antifungal phytoalexin biosynthesis (Swaminathan *et al*. 2009; Wang *et al*. 2012).

A second cluster of themes concerns environmental sensing and developmental signaling. Candidates implicate phytochrome-dependent light perception, consistent with fog’s pronounced effects on light spectral composition and irradiance (Reinhardt *et al*. 2010; Ritter *et al*. 2009): LIP1 and EID1 both regulate phytochrome-dependent signaling and could shift circadian rhythms, photomorphogenesis, and flowering time in fog-exposed plants (Hajdu *et al*. 2024; Kevei *et al*. 2007; Müller *et al*. 2018). Developmental timing candidates point more broadly to hormone and developmental regulation: AGL24 regulates the vegetative-to-reproductive transition in response to photoperiod in chrysanthemum and *Arabidopsis* (Torti & Fornara 2012; Liu *et al*. 2024), suggesting that fog may interact with light quality to modulate flowering phenology. This functional profile bears striking resemblance to the genomic basis of *Arabidopsis thaliana* adaptation to the humid, foggy precipitation regime of the Cape Verde Islands (Elfarargi *et al*. 2023), suggesting that convergent selective pressures across fog-influenced systems may act on a shared set of physiological pathways.

### Suppressed recombination at the centromere may amplify fog-adaptive selection

The centromere-adjacent fog-associated haplotype raises the question of whether its population-genetic properties reflect organismal-level selection, chromosome-level conflict, or both. Centromeres can act as selfish genetic elements, competing for preferential transmission through asymmetric female meiosis in ways that suppress recombination in adjacent regions (Zhang *et al*. 2026), precisely the architecture we observe. Our results rule out sweep-like meiotic drive, but balanced drive—in which a costly driver persists at intermediate, locally variable frequencies, as at the *Mimulus guttatus* D locus (Fishman & Saunders 2008)—cannot be excluded and testing would require transmission data from controlled crosses.

Whatever its cause, the centromeric location may itself be functionally significant: pericentromeric low-recombination regions can maintain linked gene complexes and behave as supergenes (Zhang *et al*. 2026), and large non-recombining haploblocks play a critical role in ecotype formation in wild sunflowers, where they associate with phenotypic and environmental variation consistent with local adaptation (Todesco *et al*. 2020). Suppressed recombination means that any one fog-adaptive allele among the 18 genes in this block could drag the entire haplotype toward higher frequency in foggier environments, making the centromeric architecture a potentially powerful vehicle for local adaptation regardless of whether drive is also occurring.

### Plant form tracks moisture, with potential demographic consequences

The wider growth form in foggier conditions is consistent with moisture-mediated release of auxin-driven apical dominance, with lateral branching being the more resource-responsive axis of growth— supported by reduced branching under drought in another tarweed, *Madia sativa* (González *et al*. 2008). Plants with a lower height:width ratio are more heavily branched and produce more capitula (**Fig. 4A**). If per-capitulum resource investment declines as capitula number rises, disk floret counts would follow, consistent with disk floret number being the more developmentally variable quantity (McEvoy 1984). This would also be consistent with resource-based sex allocation: disk florets are largely staminate and energetically cheaper, and plants may increase their proportional investment in male function under water stress (Tanowitz *et al*. 1987; Charnov 1982; Galen 1999). Whether these patterns reflect plasticity or genetic adaptation remains unresolved, but candidate genes from the GxE analysis are consistent with both phenotypic relationships. CNIH4, whose best-characterized role is localizing auxin transporters to the plasma membrane, and SCPL13, a regulator of cell elongation and vegetative growth, are plausible genomic links to the height:width relationship, while AP2/ERF-ANT (AINTEGUMENTA), which regulates floral organ growth, is the most direct candidate for the floret ratio shift (Wudick *et al*. 2018; Long *et al*. 2026; Elliott et al. 1996).

The broader architecture and fewer disk florets we observe in moister environments have direct ecological consequences: plant height and ray floret number promote pollinator visitation in insect-pollinated Asteraceae (Klecka *et al*. 2018), which should be especially consequential for seed set in the self-incompatible Gaviota tarplant (Tanowitz 1982; Amarilla *et al*. 2025). Ray and disk achenes also differ substantially in germination timing, dispersal mode, and seed-bank dynamics in *Deinandra* (Tanowitz *et al*. 1987), so shifts in floret composition could cascade through population demography and life history.

### Conclusion

Gaviota tarplant populations are shaped by fog and wind at multiple scales, with a coherent polygenic genomic signature of fog adaptation, morphological responses that track the same gradients, and evidence of environmental filtering during an extreme drought year. The convergence of these signals—and their parallel with *Arabidopsis* adaptation in a fog-dependent island ecosystem— suggests that the genomics of fog adaptation may be a tractable and biologically rich frontier across the world’s fog-influenced plant communities.

## Methods

### Genetic sampling, sequencing, and pre-processing

We studied the Gaviota tarplant, *Deinandra increscens* (H.M. Hall ex D.D. Keck) B.G. Baldwin subsp. *villosa* (Tanowitz) B.G. Baldwin (Madiinae, Asteraceae; Tanowitz 1982; Baldwin 1999). A subspecies of the grassland tarweed (*Deinandra increscens*), Gaviota tarplant occurs in coastal California across portions of both western Santa Barbara and San Luis Obispo counties (**Fig. S15**; Baldwin 2000). Tissue collections were made from plants in the field in May and June of 2020, 2021, and 2022 (USFWS permit no. TE-094893, CDFW permit no. 2081(a)-16-015RP). In the first year, collections were made from six occurrence localities (**Fig. 1A**): Lion’s Head (*n* = 24) and Point Arguello (*n* = 20), on Vandenberg Space Force Base; Jalama (*n* = 4); Point Conception (*n* = 22); an area between Tranquillon Mountain and Sudden Peak (hereafter, “Tranquillon Mountain”; *n* = 253); and Gaviota (*n* = 23). In the second and third years, collections were made only from Tranquillon Mountain (*n*_year1_ = 246; *n*_year2_ = 250). At each site, fresh leaf tissue from plants a minimum of five meters apart was collected onto silica gel desiccant, then accessioned to the Tissue Bank at Santa Barbara Botanic Garden.

DNA extraction followed a tarplant-specific modification (Baldwin & Wessa 2000) of a procedure using lyophilized tissue and cetyl trimethylammonium bromide (CTAB) buffer (Doyle & Doyle 1987). Remaining tissue was archived in the Tissue Bank at the Santa Barbara Botanic Garden. Whole-genome shotgun libraries were prepared for shotgun sequencing first by fragmenting DNA enzymatically with 8 min of NEBnext DNA Fragmentase treatment (New England Biolabs, Inc., Ipswich, Massachusetts), size-selective bead cleanup with PCR HiPrep beads (MagBio Inc., Gaithersburg, Maryland) at a 1:1 ratio, and then using the Spotlight protocol (Kapp, Green, & Shapiro 2021). Most libraries were dual-unique indexed to minimize the risk of index hopping; a subset used single-unique indices instead, but these were kept on separate sequencing lanes so that index hopping was not a concern. Illumina TruSeq indexes were used. Libraries were sequenced on an Illumina NovaSeq X plus at the Technology Center for Genomics & Bioinformatics at UCLA to a target depth of coverage of 1-4x, with a subset representative of sampling localities and years sequenced to a target depth of 30x.

Reads were trimmed of adapters, low-quality bases, and too-short sequences using Trimmomatic (v0.39; Bolger, Lohse, & Usadel 2014), then mapped with the Burrows-Wheeler Aligner BWA-MEM algorithm (v0.7.18-r1243-dirty; Li & Durbin 2009) to the Gaviota tarplant reference genome (McEvoy *et al*. 2024; diploid, 24 chromosomes, ∼1.7–2.3 Gb). The resulting BAM files were merged for samples sequenced across different lanes, and duplicates were marked in Picard (v2.26.2; http://broadinstitute.github.io/picard).

To identify organellar scaffolds, we aligned all scaffolds against published chloroplast and mitochondrial genomes of two other Asteraceae species, *Helianthus annuus* (mitogenome KF815390.1; plastome NC_007977.1) and *Arctium tomentosum* (mitogenome NC_058643.1; plastome MZ416781.1), using NUCmer (MUMmer v4.0.0; Marçais *et al*. 2018). Scaffolds were classified as plastid or mitochondrial when alignments of ≥ 500 bp to that organelle’s genomes covered ≥ 20% of the scaffold and the scaffold’s sequencing depth was ≥ 6× that of the chromosome-scale nuclear scaffolds; nine borderline scaffolds were assigned manually. This identified 49 plastid and 7 mitochondrial scaffolds, which were excluded from all autosomal analyses. Plastid scaffolds had higher sequencing depth relative to the nuclear genome than mitochondrial scaffolds (medians of ∼20× vs. ∼12×), consistent with plant cells carrying more copies of plastids than of mitochondria.

Merged BAM files and a soft-masked genome (McEvoy *et al*. 2024) were used as input in ANGSD (v0.940-dirty; Korneliussen & Moltke 2015) to calculate genotype likelihoods for a set of diallelic variable sites where at least 300 individuals had at least one read, total sequencing depth was between half and 2.5x the sum of the average autosomal sequencing depth of individuals (4,623x - 23,116x), and the SNP p-value was less than 1 × 10⁻⁷, leaving 47,527,450 sites. Genotypes were only called for these loci when individual sequencing depth was between a depth of 1 and twice the maximum individual depth of 60x (120x). The likelihood ratio that each of those sites was mis-mapped was calculated in ngsParalog (v1.3.4; Linderoth 2018) and sites with a Bonferroni-corrected *p*-value of less than 0.05 were discarded, leaving 36,767,362 sites. Finally, sites were filtered down to a set of 7,940,645 autosomal sites with a minor allele frequency (MAF) of >5%. For analyses that required linkage disequilibrium (LD)-filtered data, LD statistics were calculated for pairs of MAF-filtered sites in ngsLD using Tranquillon Mountain individuals sequenced to >4x depth of sequencing coverage (v1.2.1; Fox et al. 2019) and SNPs were discarded using prune_graph v0.3.5 (https://github.com/fgvieira/prune_graph) if they occurred within 50,000bp of each other and exceeded an r2 of 0.4, leaving a set of 2,221,147 LD-filtered, autosomal sites.

A set of invariant sites was also generated in ANGSD across the full sample set using standard mapping-quality/base-quality/depth filters but no minor-allele-frequency, SNP p-value, or HWE filter. Monomorphic (allele frequency < 0.001) were combined with variant sites retained through the primary SNP-calling, paralog-filtered pipeline. This all-sites set was used downstream in all SAF-based analyses (Tajima’s D, estimated effective population size, heterozygosity, etc).

### Range-wide structure and genetic diversity

Genetic structure was assessed across sampling localities using genotype likelihoods for LD-filtered sites in principal component (PCA), and admixture. For the range-wide analysis, a covariance matrix was computed for the 170 samples taken in 2020 with autosomal coverage of greater than or equal to 3x in PCAngsd (v1.36.0; Meisner, Albrechtsen, & Hanghøj 2021), the covariance matrix was decomposed, and the number of PCs retained was chosen from a scree plot. Next, an analysis of ancestral components was conducted for all 346 of the 2020 samples using a maximum likelihood approach in OHANA (v0.1.7666.41124; Cheng, Mailund, & Nielsen 2017). OHANA was run with unique seeds ten times each for two through eight ancestral components (Ks), with a minimum of 500 iterations per run. K was chosen by maximizing normalized mutual information (NMI) between each K’s hard-clustered ancestry assignments and known field-population identity; this identified *K* = 4 as the best-supported number of ancestral components, although examining a broader range of *K* values continued to reveal biologically meaningful structure among localities until *K* = 7. Genetic structure was assessed within the Tranquillon Mountain locality using a similar method; there, normalized mutual information between inferred ancestry components and plant stands increased steadily up to *K* = 6, with no meaningful improvement at higher *K*. OHANA was run in the same way on all 736 individuals, with NMI-based K selection instead being based on concordance of individuals’ assignment to ancestry components and to stands (spatial clusters) of plants. PCA was run on the subset of 250 individuals of greater than or equal to 5x autosomal coverage.

Nucleotide diversity (π) within, effective population sizes (N_e_) of, and differentiation (*F_ST_*) between populations were calculated using ANGSD (Korneliussen & Moltke 2014). The full set of variable and invariable sites was used to build the site allele frequency spectrum (SFS) for each population. For each population, the folded SFS was normalized across analyzed sites (Schmidt *et al*. 2021). For each minor allele frequency class p, the contribution to heterozygosity was calculated as 2*p*(1−*p*), weighted by the proportion of genomic sites represented by that frequency class, giving genome-wide nucleotide diversity (*π*). We estimated N_e_ using the equation θ_W_ = 4Neμ, where Watterson’s theta (*θ_W_*) was estimated for each population from the folded SFS with realSFS saf2theta and thetaStat do_stat in ANGSD v0.941, and the per-site mutation rate (*μ*) was taken from rates used for *Helianthus annuus* and *H. petiolaris* (Strasburg & Rieseberg 2008) as ∼1 × 10⁻⁸ per generation. Weighted genome-wide *F_ST_* was calculated using the realSFS fst framework in ANGSD. We tested for isolation by distance using a Mantel test comparing pairwise linearized *F_ST_* [*F_ST_*/(*1−F_ST_*); Rousset 1997] to the natural log of geographic distance between population centroids, assessing significance with an exact one-sided permutation test. Three of the four Jalama individuals had low sequencing coverage (0.9–2.9×, versus ≥ 4× in all other populations), which can bias F_ST_ between populations sequenced at different depths. To assess this, we re-estimated all pairwise F_ST_ after downsampling every other population to four individuals coverage-matched to Jalama’s (each within a factor of 1.5 of one Jalama individual’s coverage; three replicate draws), computing SAFs and F_ST_ exactly as above. Jalama’s excess differentiation relative to the other populations disappeared under coverage matching, indicating that its full-sample F_ST_ values were inflated by the coverage mismatch. We therefore excluded Jalama from the primary test (five populations), and repeated the test with all six populations using both the full-sample and the coverage-matched F_ST_ estimates. Given the small number of populations, these tests have low power, and a modest true IBD signal could go undetected.

Using a similar workflow, an SAF was built for each individual sample, and used to calculate individual *π* and pairwise *F_ST_* between individuals. To ease computational burden, we restricted these analyses to the 23 largest scaffolds (82% of the genome length). Because diversity estimates from low-coverage sequencing can be downwardly biased by allelic dropout, we modeled the relationship between mean sequencing coverage and individual diversity using a self-starting asymptotic exponential regression, which provided a substantially better fit than a log-linear model (*ΔAIC* = 79). Coverage-corrected diversity values were obtained as the residuals from this model and were used in all subsequent analyses. We tested for differences in residual heterozygosity among populations using a Kruskal-Wallis test followed by pairwise Dunn’s tests with Bonferroni correction. Folded two-dimensional SFSs were estimated from the per-individual SAFs, and weighted genome-wide *F_ST_* was calculated using realSFS *F_ST_* as previously. To test for continuous fine-scale isolation by distance within the most densely and precisely sampled population, we ran a Mantel test restricted to Tranquillon Mountain individuals only (*n* = 276). As in the population-level test, we compared linearized *F_ST_* [*F_ST_*/(1−*F_ST_*)to *ln(geographic distance)*, with significance assessed by 9,999 permutations.

Inbreeding coefficient was calculated using 500,000 subsampled LD-filtered variable sites in ngsF (v1.2.0-STD; Vieira *et al*., 2013). In a first step, the approximated estimated maximization algorithm was used with a random seed and epsilon threshold of 1×10⁻⁹, the output of which was fed as an initial value into the full algorithm. This procedure was run one hundred times, and the run producing the best final log likelihood value was retained. Inbreeding estimates are unreliable below ∼7x autosomal coverage due to insufficient power to detect heterozygotes, so we retained only samples with ≥7x coverage for inbreeding analyses. To compare inbreeding coefficients between localities, we used the same Kruskal-Wallis/Dunn’s-test approach as above, since inbreeding coefficients were also non-normally distributed and heteroscedastic.

We identified runs of homozygosity (ROH) using ROHan v1.0.5 (Renaud *et al*. 2019), which combines an initial MCMC step (--nohmm) to estimate genome-wide heterozygosity with a subsequent HMM step (--rohmu 2e-3, following calibration to the low-coverage non-human range recommended by Taylor, Manseau, & Wilson 2026) to call ROH tracts per individual. For each sample, we summed the length of all ROH tracts ≥1 Mb and expressed this as a proportion of the autosomal genome (scaffolds ≥1 Mb) to give a per-individual estimate of ROH burden reflecting recent inbreeding. The 1 Mb threshold was adopted from Ceballos *et al*. (2018), which establishes it as the conventional cutoff separating recent inbreeding from background homozygosity in wild populations: under a standard ∼1 cM/Mb recombination rate (used here in the absence of a species-specific linkage map), a 1 Mb homozygous tract corresponds to a common ancestor within roughly the last 50 generations, whereas shorter tracts reflect deeper population history (Taylor Manseau, & Wilson 2026). To test whether ROH burden differed among the three sampling years (2020–2022) at the Tranquillon Mountain site, we used a Kruskal-Wallis rank sum test followed by Dunn’s post hoc test with Bonferroni correction for pairwise year comparisons, and additionally tested the drought year (2021) against the two non-drought years combined with a Wilcoxon rank sum test.

To visualize spatial patterns of gene flow within the Tranquillon Mountain locality that may be increasing or reducing gene flow, we then computed a Fast Estimated Effective Migration Surfaces (v1.0.0 FEEMS; Marcus *et al*. 2021). We performed genotype-likelihood-based imputation on Tranquillon Mountain individuals, then sites were retained where dosage *R^2^* (a measure of imputation quality) exceeded 0.8. Genotypes were scaled prior to model fitting. The regularization parameter lambda, which controls the smoothness of the estimated migration surface, was selected by leave-one-out cross-validation (2,000 candidate values, log-spaced from 10⁻⁶ to 10⁴), and the model was re-fit using the optimal lambda to produce the final effective migration surface. This procedure was run once for all Tranquillon Mountain samples from each of the 2020, 2021, and 2022 cohorts.

### Wind connectivity analysis

To characterize the potential for wind-mediated gene flow, we built a flowering-season wind climatology representing conditions before the Strauss Wind Farm began operating in 2023: June–October of 2015–2022, the pre-turbine years for which data are available, sampled every 6 h. Wind came from analyses of NOAA’s 3-km High-Resolution Rapid Refresh model (HRRR; Dowell *et al*. 2022), with HRRR’s grid-relative wind components rotated to true north. Of the 4,896 timesteps, 4,741 (97%) were available in the HRRR archive; weighting to correct for uneven gaps among months changed no grid cell’s wind-direction frequencies by more than 0.5 percentage points. Because HRRR may not resolve the local wind acceleration and channeling expected at a coastal cape (Seto et al. 2025), we also downscaled each timestep with WindNinja (Wagenbrenner *et al*. 2016), using USGS 3DEP elevation and LANDFIRE 2016 fuel and canopy layers (LANDFIRE 2016; Rollins 2009) at 30 m resolution, with diurnal slope flows and non-neutral atmospheric stability. Validated against NOAA National Data Buoy Center station PTGC1 at Point Arguello (2015–2022, n = 4,504 timesteps; the only one of three nearby stations with observations from these years), downscaling at a 460 m mesh did not improve accuracy (wind speed mean absolute error 2.47 m s⁻¹ for HRRR vs. 3.04 m s⁻¹ for WindNinja; wind direction 19° vs. 24°). We therefore used HRRR among populations, which span several HRRR cells, and WindNinja only within Tranquillon Mountain, the largest locality in terms of census population size and location of the new wind farm. There, all sampled plants fall within roughly one HRRR cell and only downscaling can resolve terrain effects among them. Within Tranquillon Mountain, we ran WindNinja at a 100 m mesh. To assess sensitivity to these choices, we also ran a terrain-only configuration (10 m 3DEP elevation, 460 m mesh, uniform grass cover, no diurnal or stability effects) and versions adding each of the four refinements (100 m mesh, LANDFIRE vegetation, diurnal flows, non-neutral stability) separately.

Wind connectivity was modeled with the windscape R package (Kling 2026; Kling & Ackerly 2021), which summarizes each wind time series as a wind rose of directional conductance between neighboring grid cells and computes landscape wind connectivity between sites. We calculated two metrics of connectivity between sites: least-cost path (LCP) wind travel times and random walk (RW, using a range of airborne half-lives of 0.5, 3 and 24 h) deposition probabilities. Following Kling & Ackerly (2021), conductance over water (grid cells less than half land in the 3DEP elevation model) was down-weighted by 90%, wind flow from one site to another was defined as the inverse of travel time, and wind connectivity between two sites as the natural log of the mean of the flows in the two directions. Among populations, we tested whether pairs linked by weaker wind flow are more genetically differentiated using partial Mantel tests (two-sided, 9,999 permutations) relating pairwise linearized F_ST_ to wind connectivity while conditioning on the natural log of geographic distance, on environmental distance (Euclidean distance across z-scored elevation, 30-year precipitation of the warmest quarter, and 30-year fog frequency), or on both. In all population-level F_ST_ tests, Jalama was excluded from the primary analysis because its F_ST_ values were inflated by low coverage, and all six populations were analyzed as a sensitivity check. We tested the directional prediction that downwind populations should be more genetically diverse (Kling & Ackerly 2021) by correlating, for each pair of populations, the log ratio of their nucleotide diversity (*π*) with the log ratio of wind flow into versus out of each population (two-sided Mantel test, 9,999 permutations), using both LCP and RW connectivity.

Within Tranquillon Mountain, we repeated the isolation test using pairwise F_ST_ between individuals, which required hard-called genotypes and was therefore restricted to individuals with ≥ 4× coverage (*n* = 276). Tests conditioned on the natural log of geographic distance and on environmental distance among individuals, as above, and used 99,999 permutations because p-values were close to 0.05. The same tests were applied to wind connectivity from each WindNinja sensitivity configuration. To test directionality, we estimated asymmetric gene flow among all 736 Tranquillon Mountain individuals, pooled across years, with FRAME (Shen & Novembre 2026), using the same genotype-likelihood-imputed genotypes and spatial grid as the FEEMS analysis. FRAME’s smoothing parameter (*λ*) was selected by 10-fold cross-validation (*λ* = 0.00886); because cross-validation error varied little across values, we repeated the analysis at three more strongly smoothed values with near-minimal error (*λ* = 0.0379, 0.336 and 2.98). We then tested the asymmetry prediction of Kling & Ackerly (2021): for each of the 122 grid edges touching a node occupied by sampled plants, we correlated the log ratio of forward gene flow in the two directions with the log ratio of wind flow in the same two directions (*Pearson r*), using the 100 m WindNinja wind; all 337 edges were analyzed as a sensitivity check. Significance was assessed by rotating the wind field through 200 evenly spaced angles, which preserves its spatial and temporal structure while changing its direction; the two-sided p-value was the proportion of rotations yielding a correlation at least as strong (in absolute value) as that observed. All within-Tranquillon tests were based on LCP; the random-walk alternative was omitted because pairwise computation scales poorly to 736 individuals and because the half-life decay parameter has little effect over the short inter-plant distances on Tranquillon Mountain..

### Environmental variable selection

We started with a set of environmental variables of potential interest to the local adaptation of the Gaviota tarplant, broadly composed of variables characterizing soil physical properties, topography, bioclimate, climate water deficit, and fog regime. Variables were excluded from this candidate set if they were categorical or near-invariant, or if the native spatial resolution of the dataset was less than 1 km^2^. For similar variables, we compared their definitions and retained the one we considered more ecologically-relevant to the tarplant. Since aspect could not be meaningfully assessed with Pearson/Spearman correlation because of its wrap-around discontinuity, it was composed via unit-circle projection into two linear components, northness (cos) and eastness (sin). Once we had a set of non-redundant candidate variables, we subsampled environmental data so that all plant sampling localities were represented equally. Using this normalized dataset, we pruned the candidate variable set for collinear variables (*rho* ≥ 0.55) using pairwise Spearman correlations, favoring the variable with finer spatial resolution or more direct ecological interpretation.

The final variable set (**Fig. S16**) comprised four variables spanning fog exposure, summer precipitation, precipitation seasonality, and climatic water deficit, each averaged over a common 2000-2025 reference window and summarized via a centered and scaled principal components analysis to characterize the dominant axes of environmental variation among sampled populations. Fog exposure was derived from Fog and Low-lying Cloud Cover (FLCC; Werner *et al*. 2022), the frequency of daily summertime marine fog and low cloud presence detected within each pixel from MODIS cloud flags, expressed as the mean number of days per month with fog/low cloud presence during the fog season (June-September). Climatic water deficit (CWD), a monthly measure of plant-relevant water balance integrating water supply against atmospheric demand, was derived from the California Basin Characterization Model, a physically based hydrologic model (Flint & Flint 2014). Summer precipitation and precipitation seasonality were derived from PRISM gridded climate data (Daly et al. 2008): summer precipitation as the multi-decadal mean of total precipitation summed over June-September, and precipitation seasonality as the multi-decadal mean of the coefficient of variation among each year’s monthly precipitation. For each specimen, summary metrics representing two temporal scales were computed. One was a germination-year mean value, and the other a rolling 20-year mean ending in, and including, that specimen’s own germination year. Years were always treated as “water years”, meaning October of the prior year through September of that year.

### Genomewide signatures of local adaptation

For selection analyses, the imputed genotype file computed for FEEMS was filtered to remove individuals with autosomal coverage less than four (leaving *n* = 336). Associations between genotype and the four retained environmental variables were tested using GEMMA v0.98.3 (Zhou & Stephens 2012). We used MAF-filtered biallelic SNPs, individuals with at least 4x depth of sequencing coverage, and a univariate linear mixed model (-lmm 2, likelihood-ratio test). We included an explicit intercept, the first five principal components of genome-wide genotype covariance (estimated in PCAngsd from LD-pruned sites), and mean per-individual sequencing depth as covariates, and also controlled for a genome-wide centered kinship matrix computed in GEMMA from the same unpruned SNP set. Test-statistic calibration was evaluated using the genomic control statistic *λ* (median observed *χ²* / median expected *χ²* under the null), computed from the LRT p-values. Genomic control was applied if *λ* exceeded 1.05. SNPs with likelihood-ratio *p*-values below a Bonferroni threshold (*α* = 0.05 / total number of SNPs) were considered genome-wide significant. Bonferroni-significant SNPs were further classified as belonging to a co-localized peak if they fell within a 40 kb window (sliding in 20 kb steps) containing at least three significant SNPs.

The associated gene(s) and predicted effect type (genic, bounded upstream/downstream, or intergenic) for each SNP were annotated using SNPeff v5.2e (Cingolani *et al*. 2012) and a custom-built *Deinandra increscens* subsp. *villosa* database. An intergenic-only SNP retained its nearest-gene association only if it was in linkage disequilibrium (*r²* ≥ 0.5) with a variant inside that gene’s body (based on a precomputed genome-wide LD scan); genic and distance-bounded upstream/downstream calls were retained unconditionally. To confirm that candidate genes nominated through genotype-by-environment association are actively expressed in *D. increscens* subsp. *villosa*, reads from eight RNA-seq libraries spanning root, shoot, and leaf tissue from a single Tranquillon Mountain population (McEvoy *et al*. 2024) were quantified against the *D. increscens* transcriptome, extracted from the masked reference genome using the longest-isoform GFF annotation. Expression was summarized as mean transcripts per million (TPM) per gene.

To test whether residual departure from the null p-value distribution after genomic control reflected genuine genotype-phenotype covariation rather than test-statistic miscalibration, we performed a permutation test for each environmental variable in which phenotype values were randomly permuted 100 times across individuals while genotypes, the kinship matrix, and covariates were held fixed. Variables were not pooled into a shared null, since their marginal distributions differ substantially in shape. For these tests, we used a genome-wide SNP subset for computational tractability, sized per variable to ensure adequate representation of that variable’s significant loci. λ and Bonferroni-hit counts from the observed and permuted data were compared to derive empirical p-values, calculated as the proportion of permutations at least as extreme as observed, with a +1 correction to numerator and denominator to avoid zero p-values (Davison & Hinkley 1997).

To test whether the observed gene-set overlaps between environmental variables exceeded chance expectation, we performed pairwise hypergeometric tests treating each variable’s candidate gene set as a sample drawn without replacement from the genome-wide complement of annotated genes. For each pair of variables, we calculated the probability of observing at least as many shared genes as were actually observed, given the two sets’ sizes and the total gene universe, using the hypergeometric survival function), with the expected overlap under independence computed as the product of the two set sizes divided by the gene universe size.

### Gene ontology enrichment & tissue-level expression of candidate genes

Gene Ontology (GO) annotations for the *Deinandra* gene set were compiled from five independent sources and combined by union. First, orthology-based functional annotation was assigned with eggNOG-mapper v2.1.15 (Cantalapiedra *et al*. 2021) run against the translated coding sequences. Second and third, for genes receiving no GO terms from eggNOG-mapper, and as a supplementary check for genes that did, GO terms were additionally harvested from the NCBI Gene2GO database (Ashburner *et al*., 2000; Gene Ontology Consortium, 2026) via each gene’s best BLASTp hit. Fourth, orthology-based transfer from *Arabidopsis thaliana* was performed as follows: coding sequences were translated to protein, and orthology between *Deinandra* and the *A. thaliana* TAIR10 reference proteome (Ensembl Plants) was inferred with OrthoFinder v3.1.5 (Emms & Kelly 2019). Fifth, GO terms were transferred in the same way from sunflower (*Helianthus annuus*, HanXRQr2.0-SUNRISE), whose proteome was included in the same OrthoFinder analysis, using its non-electronic (non-IEA) annotations from Ensembl Plants BioMart. GO terms were transferred only for genes in strict single-copy orthogroups (i.e., orthogroups containing exactly one *Deinandra* and one gene from the source species), using direct (non-electronic, evidence code other than IEA/ND) TAIR annotations. GO terms from all four sources were propagated to their ancestor terms under the true-path rule (is_a/part_of relations) using a graph parsed directly from a current release of the GO basic ontology (go-basic.obo), rather than cached Bioconductor annotation packages, which lag current GO releases.

GO enrichment was tested using continuous association evidence. For each environmental variable, genomic-control-corrected per-SNP GEMMA p-values were combined into a single gene-level statistic using the Cauchy combination test (ACAT; Liu & Xie 2020). Genes were ranked by their ACAT-combined−log10(p) and tested for GO term enrichment along this ranking using fgsea v1.32.4 (Korotkevich *et al*. 2021) with an unsigned (single-tailed) scoring scheme, appropriate because the ranking statistic has no natural directionality. Non-plant-plausible and housekeeping terms were excluded from the tested category set prior to running fgsea, preserving the correct multiple-testing denominator for BH correction across the remaining, biologically relevant categories. Because no term reached genome-wide FDR significance for any of the three environmental variables, we present nominally significant terms (*raw p* < 0.05, and *p* < 0.005 for the summary figures) with redundant GO terms collapsed within each ontology (BP/MF/CC) using rrvgo v1.18.0 (Sayols 2023; Wang semantic similarity, threshold 0.7), retaining one representative term per cluster. To assess whether nominal signal reflected more than chance, we additionally tested for convergence across the three independent GEMMA analyses: GO terms recurring as nominal hits with identical GO identity in ≥ 2 variables; GO terms with distinct but highly similar identity (*pairwise GO semantic similarity* ≥ 0.8, Wang method, *A. thaliana* annotation via GOSemSim) in ≥ 2 variables, reported separately as near-miss candidates rather than treated as confirmed matches; and, separately, leading-edge candidate genes recurring across nominal terms in ≥ 2 variables.

### Genomic context and dynamics of the DEINVI_0000G fog-associated haplotype

The DEINVI_00006 fog peak sits within an unusually long block of elevated association (**Fig. 3C**), prompting us to investigate whether a structural variant underlies the signal. To follow this region in every sampled plant, we used genotype dosages imputed from genotype likelihoods for all 842 sequenced plants (Beagle r1399; sites with DR2 > 0.8). The favourable allele was defined as the allele whose dosage, after regressing out sequencing coverage, increased with long-term fog across all plants. We ran windowed PCA (Blumer, Good, & Durbin 2025; 1-Mb windows, 10-kb steps, genotype likelihoods, MAF ≥ 0.05) along DEINVI_00006, correcting window-to-window sign flips of PC1; this identified a block (∼45.3–49.3 Mb) in which plants fell into three PC1/PC2 groups with a sharp right edge at 49.1–49.4 Mb. We searched for breakpoints using short-read pairs with mates or split-read parts mapping > 20 kb apart, assigned to plants grouped by k-means on PC1/PC2 (≥ 90% of windows; MAPQ ≥ 20), testing for each pair of linked 50-kb bins whether one homozygous group carried supporting reads more often than the other (one-sided Fisher’s exact test, Bonferroni-corrected). No group-specific link joined the two edges of the block; any group-specific links were local and did not form a consistent pair of junctions. We aligned PacBio HiFi and Oxford Nanopore reads from the reference individual (McEvoy *et al*. 2024) to the assembly with minimap2 v2.24-r1122 (Li 2018) and found no inversion breakpoints: the right edge was crossed intact by 23 spanning reads, the only left-edge junction was present in 89% of plants across all groups (a fixed or assembly feature), and clipped reads clustered throughout the repeat-rich interior. To test whether LD decays normally within homokaryotype groups—as it should if the block is not an inversion—we assigned Tranquillon Mountain plants to three groups by k-means clustering of mean PC1 over the block core (46.3–49.1 Mb), subsampled each to 36 plants, and computed median *r²* between imputed dosages (within-group MAF ≥ 0.10) in distance bins of 1–300 kb, comparing the core with control regions on the same chromosome (2–8 and 55–61 Mb).

To instead explore whether the block represented a centromere, we calculated gene density and repeat-family content in 500-kb windows from the repeat-annotated genome (McEvoy *et al*. 2024). Tandem satellite arrays were identified de novo from 21-mers occurring ≥ 200 times on a scaffold, with monomer length taken as the modal spacing between successive occurrences of the most frequent k-mer. Once we had identified the centromere on DEINVI_00006, we defined two centromeric haplotypes from the outer PC1 groups of the all-locality windowed PCA, identified 649 SNPs in the centromeric core whose frequency differed by ≥ 0.8 between them, and assigned each plant a hybrid index (mean A-allele dosage / 2; AA > 0.75, BB < 0.25, otherwise AB); for the fog peak, we calculated *r²* between each plant’s mean favourable dosage over the 19 peak SNPs and the imputed dosages of all SNPs (MAF ≥ 0.02) at 42–52 Mb in the 736 Tranquillon Mountain plants, and compared the favourable haplotype’s frequency between plants homozygous for each centromeric haplotype (same group in ≥ 90% of windows). Finally, because outliers can be over-represented in low-recombination regions even under neutrality (Booker, Yeaman & Whitlock 2020), we tested whether outlier peaks are enriched near centromeres: peaks within 2 Mb of one another were merged into loci, each locus’s distance from its centromere was expressed relative to its chromosome arm length, and the mean relative distance and fraction within 5 Mb of a centromere were compared with 10,000 sets of random tested SNPs on the same scaffolds (one-sided p), for all Bonferroni-significant peaks associated with fog, precipitation seasonality, and summer precipitation on scaffolds with confident centromere calls. Because centromere-proximal loci are prone to collapsed or ambiguously mapped repeats, we tested whether this association could instead reflect a mapping artifact: For each 10-kb window along DEINVI_00006, we computed pooled read depth across all individuals with samtools depth (Li *et al*. 2009), once with no mapping-quality filter and once restricted to reads with MAPQ ≥ 20, took the proportion of depth lost under the MAPQ ≥ 20 filter as a window-level measure of ambiguous/low-confidence mapping, and compared this fraction between the GWAS peak and the rest of the genome (excluding the centromere) with a one-sided Mann-Whitney U test (*n* = 11 peak windows vs. 6,690 background windows, *U* = 30,705, p = 0.83)..

To determine whether the DEINVI_00006 centromere and the fog peak sit at a junction between ancestral chromosomes, we identified syntenic genes between *D. increscens* and sunflower (*Helianthus annuus*, HanXRQr2.0-SUNRISE), *Helianthus anomalus* (Hanom1.0, GCA_047456115.1), lettuce (*Lactuca sativa*, Lsat_Salinas_v11, GCA_002870075.4), *Arabidopsis thaliana* (TAIR10) and tomato (*Solanum lycopersicum*, SL4.0), using annotations from Ensembl Plants release 63 or NCBI (*H. anomalus*), with GENESPACE v1.3.1 (Lovell *et al*. 2022), retaining gene pairs within syntenic blocks. Lettuce, which lies outside the genome triplication shared by sunflower and *Deinandra* and so has one counterpart for each *Deinandra* region, served as the primary reference for ancestral chromosome structure, with sunflower as support. For each of the chromosome-scale scaffolds we followed, from the syntenic gene nearest each edge of the centromere call outwards, the run of genes whose partner lay on the same chromosome (allowing up to two consecutive genes with another partner), and classed the centromere as a junction between ancestral segments when the two flanking runs had different partner chromosomes, as continuous when they shared one, and as ambiguous when either run had fewer than five genes; the lettuce-based classifications did not change across alternative thresholds (0-5 stray genes; minimum runs of 3-10 genes) except that some continuous scaffolds became ambiguous at the strictest setting. To explain why GNS1 and SNF1 are adjacent in *D. increscens*, we located their syntenic orthologues in sunflower and lettuce, listed all genes lying between them, placed each intervening gene in *D. increscens* by its syntenic orthologue or, failing that, its best BLAST hit, and tested collinearity (Spearman’s rho) between lettuce and each *D. increscens* chromosome carrying syntenic genes within 3 Mb of the interval.

Because the favourable haplotype’s landscape-scale association with fog rests largely on a single locality, we tested whether it tracks fog within Tranquillon Mountain, where repeated sampling of spatially structured stands in three years (2020-2022) allows spatial and temporal effects to be separated. Each peak was summarised for each plant as the mean favourable-allele dosage across all of its retained outlier SNPs (SNPs ≤ 40 kb apart; ≥ 3 SNPs per peak), each SNP having been polarised separately to its fog-favourable allele; for DEINVI_00006, near-complete linkage among all 19 SNPs means this behaves as a single haplotype dosage, with only 17 of 736 Tranquillon Mountain plants ambiguous (plants > 0.25 from a whole number of copies were counted as ambiguous). Haplotype frequency was this dosage divided by two, estimated per locality across the range and per unit within Tranquillon Mountain. Plants (n = 736) were grouped into stands, one stand spanning several fog grid cells being split by cell, and units of < 5 plants were dropped, leaving 13 units. For the spatial test, the statistic was the plant-weighted least-squares slope of unit haplotype frequency on unit long-term fog; for the temporal test, it was the slope of each plant’s haplotype frequency on germination-year fog centred within its unit, so that only change over time within a place contributed. Significance (one-sided) was assessed by permuting long-term fog among units or germination-year fog among plants within units (10,000 permutations each); against SNPs from 8,000 random 2-kb regions elsewhere in the genome (one SNP per region), polarised in the same way and matched to the peak’s Tranquillon frequency (±0.04); and, because allele frequencies near centromeres may fluctuate more, against such SNPs drawn at a similar distance from their own centromere as the peak. The same statistics were computed for each of the other 30 fog peaks, each summarised in the same way as the mean over its own SNPs, as a locus-level comparison set sharing the same plants, years, units and polarisation.

Because the peak lies beside a centromere, a classic site of female meiotic drive, we tested three predictions of a driver spreading through populations. First, Tajima’s D should be low for the peak’s genomic setting in every population. For each locality with sufficient sample size (excluding Jalama; individuals with ≥ 4× coverage), site allele frequencies were estimated in ANGSD, the folded SFS was estimated with realSFS and used as a prior to compute per-site Watterson’s *θ* and *π* with thetaStat, and Tajima’s D was calculated in sliding 100-SNP windows. Fog outlier SNPs within 40 kb of one another were merged into peaks, and each peak was summarized as the median Tajima’s D of the windows overlapping it (115 kb for the DEINVI_00006 peak). The DEINVI_00006 peak was compared with 3,000 random 115-kb blocks lying outside the centromere call at a similar distance from their own centromere (0–1.6 Mb), and with 3,000 blocks of the same length placed anywhere on scaffolds with a confident centromere call; the other 30 fog peaks were compared with the latter. Blocks excluded DEINVI_00006 and ± 500 kb around any fog peak and required ≥ 20 windows. One-sided p-values were the proportion of blocks with a median Tajima’s D at or below the peak’s, with a +1 correction to numerator and denominator. Second, a driving centromeric haplotype should approach similar frequencies everywhere: from the hybrid-index genotypes we estimated the frequencies of centromeric haplotypes A and B per locality and, in Tranquillon Mountain, the inbreeding coefficient *F* = 1 - *H_obs_*/*H_exp_*with an exact Hardy-Weinberg test, comparing F with that computed in the same way from random genome-wide SNPs. Third, a driver spreading by gene flow should reduce differentiation at the centromere: pairwise *F_ST_* was estimated in ANGSD (realSFS) in 50-kb windows with 10-kb steps (windows with >= 100 sites), and the median *F_ST_* of windows within the DEINVI_00006 centromere call was compared with the genome-wide window distribution and with the same statistic at the other centromeres.

Because centromere drive is expected to select for rapid adaptive evolution of the centromeric histone CENH3, particularly in its N-terminal tail (Finseth *et al*. 2015, Zedek & Bureš 2016), we compared its rate of evolution with the genomic background. CENH3 was identified by tblastn of sunflower CENH3 tails against the assembly; the single hit is the 3’ end of a gene model mis-merged with an adjacent Argonaute gene, which we trimmed to the CENH3 start codon. Coding sequences were obtained for *H. annuus* (Ensembl Plants), *H. anomalus* (NCBI), *H. debilis* (NCBI), *Layia glandulosa* (miniprot model on the alternate-haplotype assembly GCA_046128875.1), lettuce (histone-fold domain only) and *A. thaliana*. Proteins were aligned with MAFFT v.7.526 (L-INS-i; Katoh *et al*. 2002) and back-translated with PAL2NAL v14.1 (Suyama, Torrents, & Bork 2006), and the tail was defined as alignment columns before the conserved histone-fold motif (R[FY][KR]PGT, located in *H. annuus*). Pairwise dN/dS was estimated with PAML v4.10.10 (Li *et al*. 2018) for the whole protein, tail and histone fold, and placed against 1,995 random *D. increscens*-*H. annuus* one-to-one syntenic orthologues processed identically. Site models (M1a vs M2a, M7 vs M8) and the branch-site model A on the *Deinandra* branch were fitted with PAML and compared by likelihood-ratio tests.

### Phenotypic responses to environmental moisture

To test whether variation in floral and vegetative traits tracks environmental moisture availability, we related two phenotypes (the proportion of florets that are ray florets and the ratio of plant height to width) in relation to fog exposure and summer precipitation, the two moisture-related variables retained from the genotype-by-environment variable set (precipitation seasonality was excluded from these models; see below). Phenotypic data were derived from herbarium specimens and field measurements collected across the range of the Gaviota tarplant (**Fig. S17**). New specimens accessioned for this project were collected from seven sites across the range of the subspecies, ranging from 12–432 m elevation, in summer of 2023 and 2024. Localities sampled included the six sites from which genetic samples were taken, as well as a seventh, between Point Conception and Gaviota (“Agujas”). Plant height (ground to the highest point of the plant, without straightening the stem) and width (widest point as seen from above, similarly without straightening) were measured in the field; plants were pressed and accessioned to the Santa Barbara Botanic Garden Clifton Smith Herbarium (SBBG). Additional SBBG herbarium specimens, collected between 1927 and 2024, extended the phenotypic record further back in time.

One capitulum from each of 156 specimens was dissected under a dissecting microscope to count ray and disk florets (used together to calculate the proportion of florets that are ray florets); for mounted historic specimens, capitula not glued to the herbarium sheet were chosen, and for all specimens we selected mature, complete capitula (based on visible anther presentation) discrete from neighboring floral heads. Rays and disks were mounted on glass slides and stored at the UC Santa Cruz Kenneth S. Norris Center for Natural History Herbarium (UCSC). We first tested whether the proportion of ray florets and height:width ratio differed among sampling localities, independent of any environmental covariate. Because this treats locality as a fixed grouping factor (rather than the random effect used in the environmental models below), it was restricted to modern (post-2000) specimens, excluding CDFW and Jalama (*n* = 6 and *n* = 4, too few for a reliable fixed-effect comparison). Neither trait was normally distributed, so we used the Kruskal-Wallis/Dunn’s-test approach, Bonferroni-corrected, where the omnibus test was significant. Ray floret count across time was supplemented with additional counts from 137 photos of specimens in the field from community science platform iNaturalist.org. It was not possible to count disk florets from photos. iNaturalist specimens included 86 specimens accessioned to SBBG for this project in 2024, as well as 51 observations without an associated physical specimen, observed between 2010 and 2026. Ray florets were counted from one capitulum per observation. Capitula were selected if the capitulum appeared to be complete, with no missing flowers, and all rays were visible. A paired Wilcoxon signed-rank test demonstrated that iNaturalist ray floret counts and dissected ray floret counts did not differ significantly (*V* = 580, *n* = 85, *p* = 0.425).

To test whether these traits respond to fog exposure and summer precipitation specifically, rather than to locality identity per se, we built generalized linear mixed models (GLMMs) with locality as a random intercept -- retaining every locality regardless of sample size, since a random effect appropriately shrinks a small locality’s estimate toward the grand mean rather than requiring exclusion. For each modern specimen, fog exposure and summer precipitation were each calculated as a 20-year rolling mean ending in (and including) the specimen’s own germination year. Precipitation seasonality was tested as a third candidate predictor but dropped after proving collinear with fog exposure in the phenotyping dataset (*Spearman rho* = −0.84 to −0.89 depending on window length). Climate water deficit was tested as a candidate predictor but did not improve model fit for either phenotype. For each of the two phenotypes we fit every additive and interactive combination of the two z-scored predictors (fog exposure, summer precipitation, climate water deficit, additive, interactive), with locality as a random intercept and an error distribution matched to the trait (beta regression for ray floret proportion; Gamma for height:width ratio), and ranked candidates by AICc. Because many models and terms were compared, results are exploratory rather than confirmatory, without correction for multiple comparisons; a model was considered well-supported only when its intercept and every fixed-effect coefficient were significant (*p* < 0.05). As a robustness check, we refit the top model for both phenotypes using each specimen’s own single germination-year value instead of the 20-year rolling mean. Height:width ratio was available only for specimens field-measured in 2024, at Lion’s Head, Point Arguello, Tranquillon Mountain, and Point Conception; ray floret proportion required a physical specimen (see above).

To test for a trend in ray floret proportion and disk floret count over time, independent of locality, we fit generalized linear mixed models (beta-regression and negative-binomial, respectively; value ∼ year + (1 | Locality)) across the full historic record (1927–2024; *n* = 162 dissected specimens), and an equivalent model for height:width ratio (Gaussian GLMM; *n* = 115 modern specimens). Trends in fog frequency (> 1 mm total, 1950–2014) and the probability of summer precipitation (1927–2024) over the historic record were assessed with non-parametric Mann-Kendall tests on annual means averaged across all sampled coordinates. Since the Werner et al. (2022) MODIS-based fog dataset only extends back to 2000, fog frequency trends were drawn from Kling (2016), which spans 1895–2014; see **Fig. S18** for comparison of the two fog metrics.

## Acknowledgements

A permit for this work was provided to Santa Barbara Botanic Garden (SBBG) by the US Fish and Wildlife Service (Recovery Permit Number ES094893). For assistance with field site access, we thank Kelly Schmoker of the California Department of Fish and Wildlife; Elizabeth Hiroyasu, Moses Katkowski, Gabriela Morales, Chelsea Nielsen, Mark Reynolds, and Laura Riege of The Nature Conservancy’s Jack and Laura Jalama Preserve; Jeffrey Carroll, Elihu Gevirtz, Richard Podolsky, Javier Venegas, of the Strauss BayWa r.e. wind farm; and Luanne Lum of Vandenberg Space Force Base. Field collections were made possible with the assistance of SBBG botanists Annie Lamont-Ayers, Caitlin Hazelquist, Jessica Spaw, Isabella Salangsang, and Sean Carson, and Isabel Rivera provided valuable assistance with laboratory work at SBBG. Phenotypic work at the Norris Center was facilitated by Gizelle Hurtado and Al Keuter. We thank Bruce Baldwin, Mark Elvin, Jake Marcon, Barry Tanowitz, and Dieter Wilken for generously sharing their knowledge of Gaviota tarplant, and Alicia Torregosa, Ryan Martin, and David Ackerly for their assistance in understanding regional environmental datasets. Thank you for computational support to the UC Santa Cruz Genomics Institute, particularly Erich Weiler; and to Rasmus Nielsen for advice with population genetic analysis. Funding for this project was provided by BayWa r.e. E.S. was supported by the University of California President’s Postdoctoral Fellowship Program and by a National Science Foundation Postdoctoral Research Fellowship in Biology. D.S. was supported by the Research Mentoring Internship program at UCSC. C.M.G. was supported by an endowment gifted to Santa Barbara Botanic Garden by Ken and Shirley Tucker.

## Notes

### Competing Interest Statement

The authors have declared no competing interest.

### Summary of Updates

Improved wind connectivity analysis using WindScape, including an updated test of directional impacts on gene flow using FRAME for asymmetrical migration estimates. Updated figure 4 to include summer precip trend map. Large-scale word count cut.

