## Supplemental Materials for "Fog adaptation and wind connectivity in an endangered California plant: a genomic baseline for wind energy development"

### Supplementary Materials

#### Contents


| | | |
|---|---|---|
| **Fig. S1** | Admixture analysis of 2020 data across Ks 2 to 8 | Page 2 |
| **Fig. S2** | Nucleotide diversity of *Deinandra increscens villosa* populations | Page 3 |
| **Fig. S3** | Admixture analysis of Tranquillon Mountain data across Ks 2 to7 | Page 4 |
| **Fig. S4** | Pairwise Fst between pairs of Tranquillon Mtn individuals | Page 5 |
| **Fig. S5** | Effective migration surfaces for Tranquillon Mountain site | Page 5 |
| **Fig. S6** | Wind connectivity and genetic differentiation at two spatial scales | Page 6 |
| **Fig. S7** | Manhattan plots comparing outlier peaks and annotated genes between fog, summer precipitation, and precipitation seasonality | Page 7 |
| **Fig. S8** | Candidate gene overlap across fog and low lying cloud cover, precipitation seasonality), and summer precipitation. | Page 7 |
| **Table S1** | Outlier genes relative to fog exposure, precipitation seasonality, and summer precipitation | Page 8-10 |
| **Fig. S9** | Enriched gene ontology terms among candidate genes associated with fog, precipitation seasonality, and summer precipitation | Page 11 |
| **Fig. S10** | Tissue expression of candidate adaptation loci | Page 12 |
| **Fig. S11** | Fog-associated outlier peaks and their position relative to the inferred centromere | Page 13 |
| **Fig. S12** | GxE outliers, structure, and synteny along chromosome DEINVI_00006 | Page 14 |
| **Fig. S13** | Frequency of the centromeric-associated haplotype in chromosome DEINVI_00006 between germination years. | Page 15 |
| **Fig. S14** | Tajima's D at the DEINVI_00006 fog peak and other fog outlier peaks, compared with random genomic blocks | Page 15 |
| **Supp. Note** | The centromeric histone CENH3. | Page 16 |
| **Table S2** | Top three models, by AIC, resulting from model comparison of how fog, summer precipitation, and their interaction might contribute to the proportion of ray florets and the height to width ratio | Page 16 |
| **Fig. S15** | Phylogenetic and geographic context for the Gaviota tarplant (*Deinandra increscens* subsp. *villosa*) | Page 17 |
| **Fig. S16** | Collinearity and PCA of retained environmental variables for genotype by association analyses | Page 17 |
| **Fig. S17** | Map of new and historic specimens used in phenotype analysis | Page 18 |
| **Fig. S18** | Comparison of the two fog metrics used in this publication | Page 18 |

**Figure S1.** Admixture analysis of 2020 data across Ks 2 to 7. Facets bin samples by locality, and within facets samples are sorted by longitude. The predominant ancestral component within a locality is colored to match the color assigned to that locality.

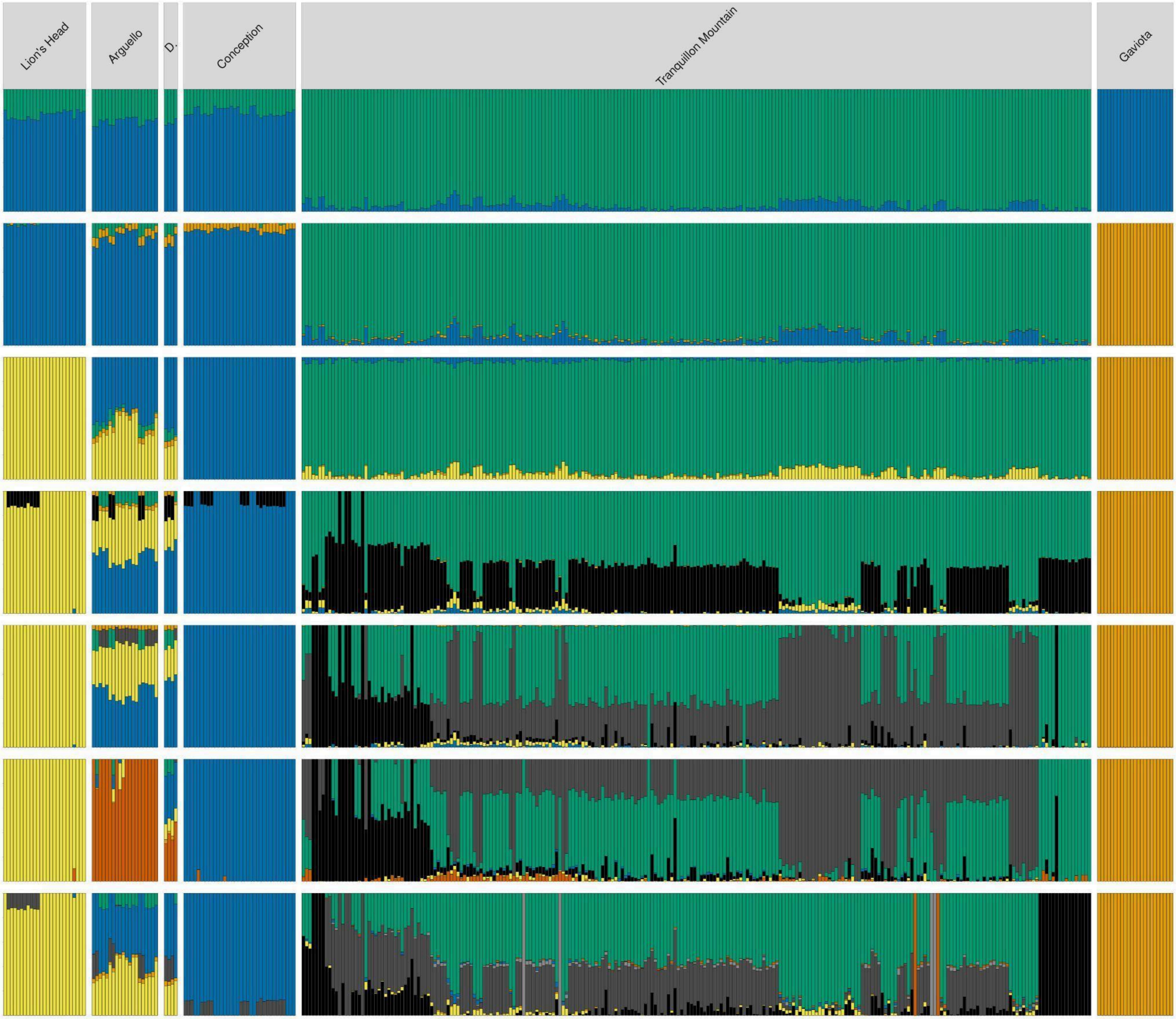

**Figure S2.** Nucleotide diversity of *Deinandra increscens villosa* populations, compared to populations of other flowering plants (Angiospermae), Conifers (Coniferae), and green algae (Chlorophyta), taken from Leffler *et al.* 2012.

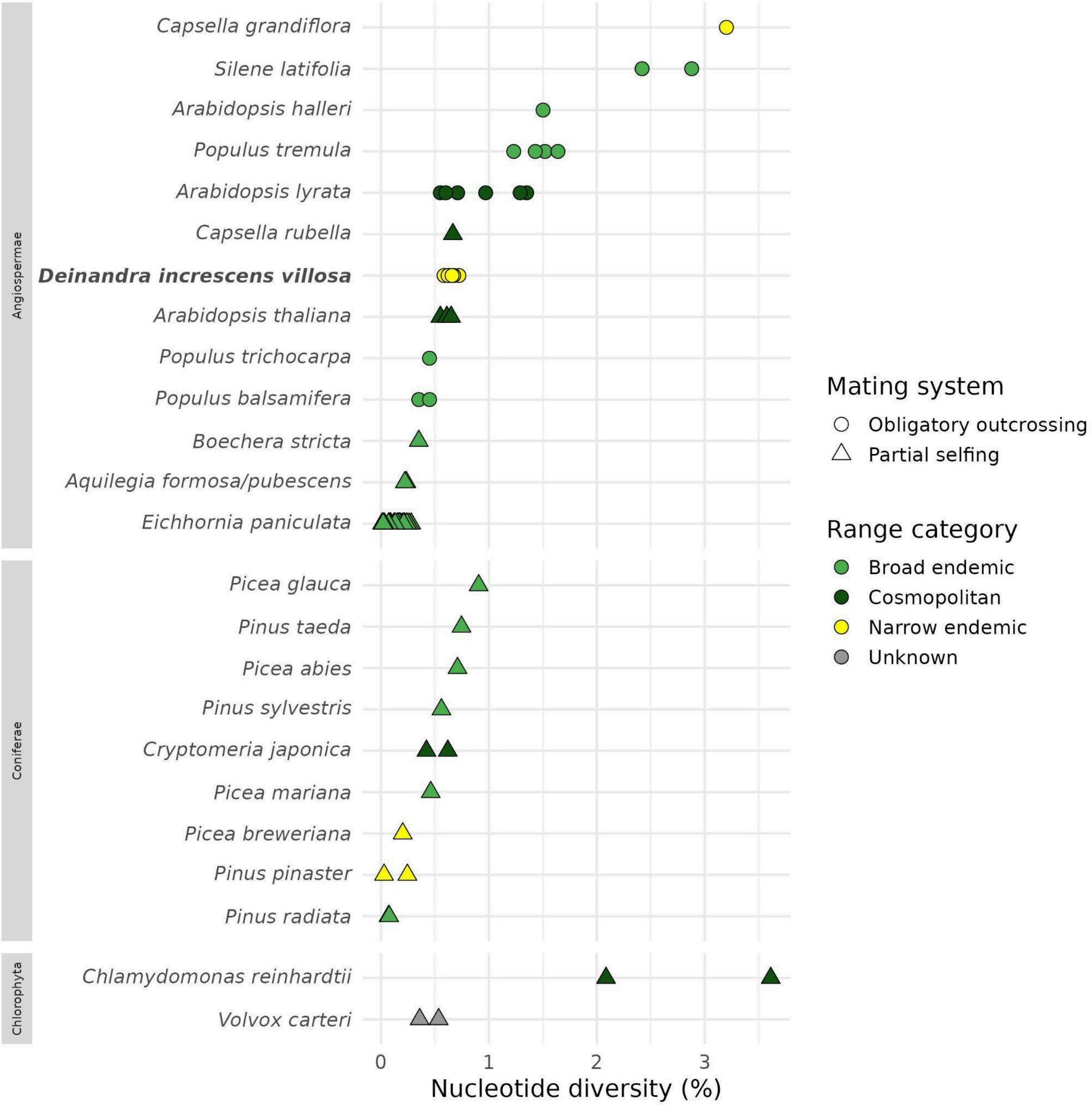

**Figure S3.** Admixture analysis of Tranquillon Mountain data across Ks 2 to 7. Samples are sorted by clusters of plants, and generally by longitude from west to east. The y-axis represents the proportional composition of individuals to the different possible number of ancestral components.

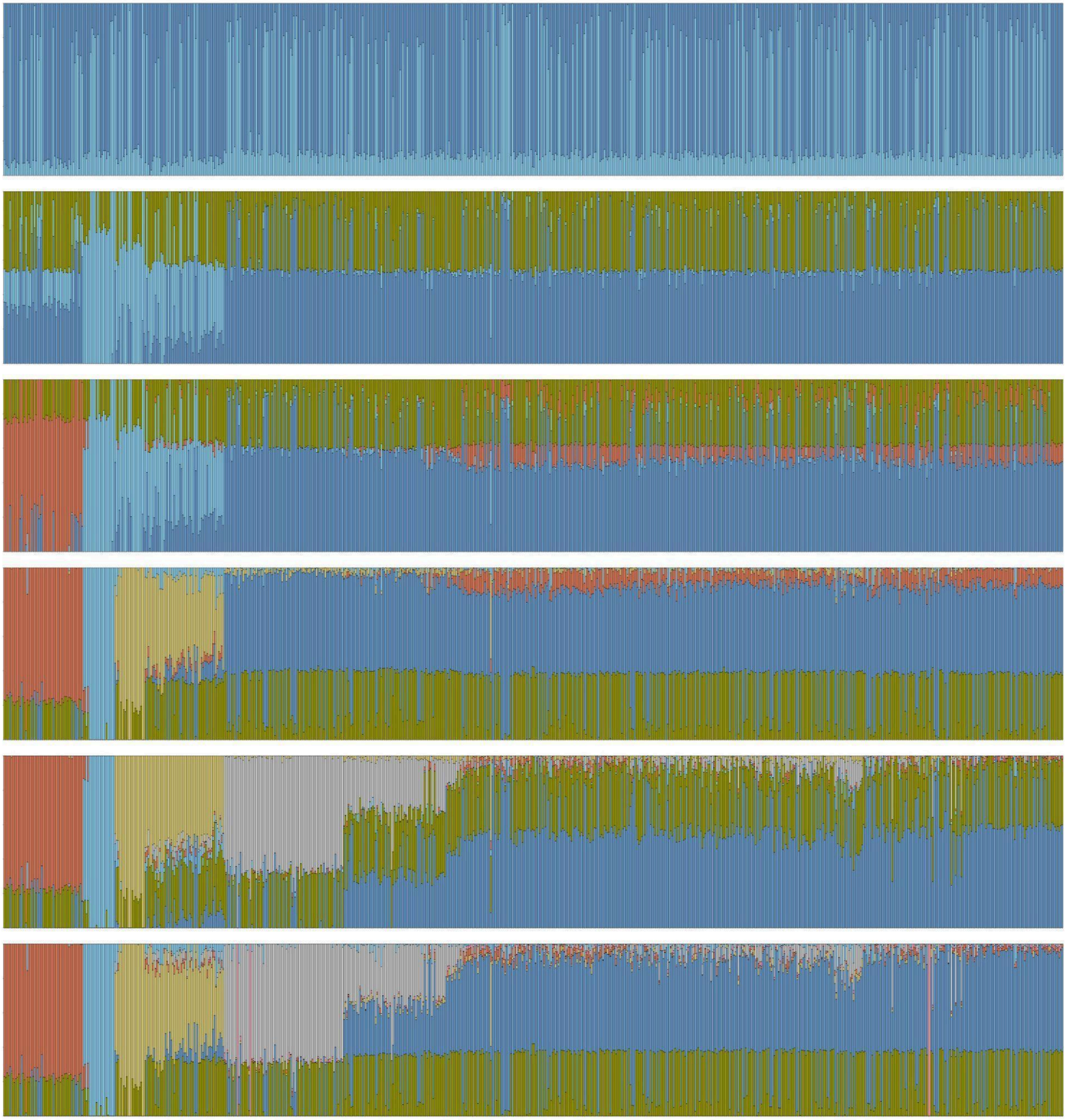

**Figure S4.** Pairwise Fst between pairs of Tranquillon Mtn individuals (green). The small effect size ($r^2 \approx$ 1.4%) suggests that geographic distance explains only a fraction of the genetic variance among individuals within this population, consistent with the weak relationship seen in the broader population-level analysis.

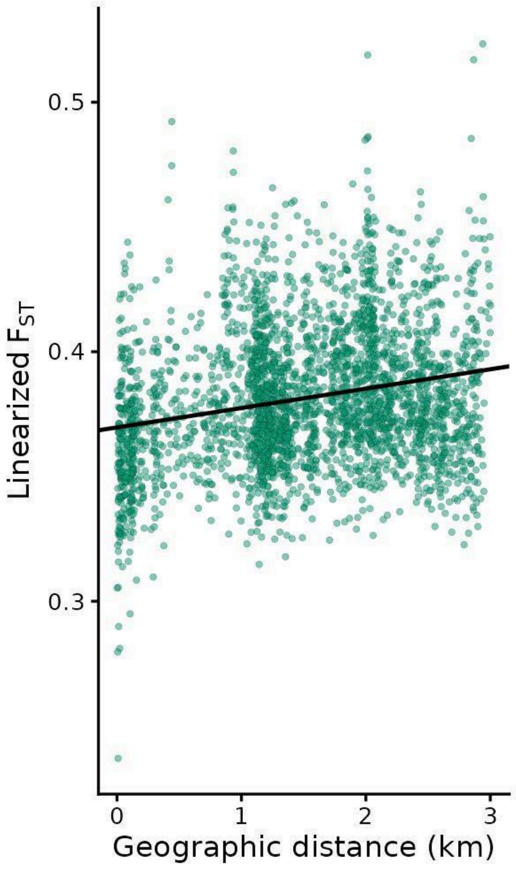


**Figure S5.** Effective migration surfaces for Tranquillon Mountain site. The panels below show surfaces for 2020 (λ = 253.1), 2021 (λ = 452.3), and 2022 (λ = 96.2) samples, calculated separately.

2020 alone 2021 alone 2022 alone

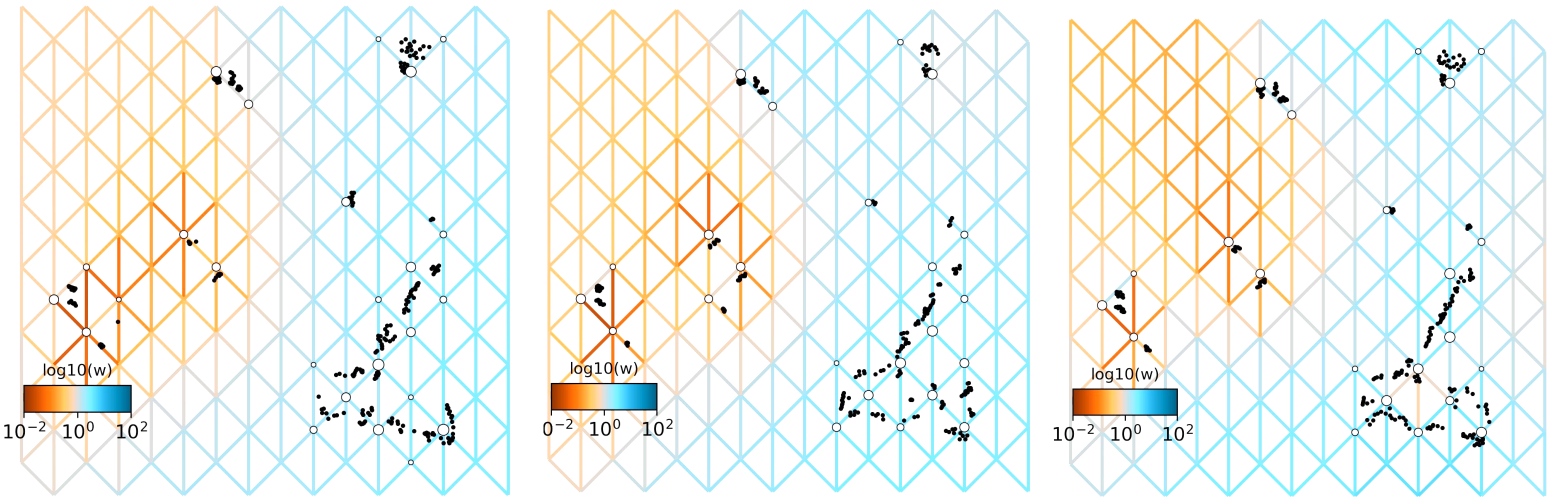

**Figure S6**. Wind connectivity and genetic differentiation at two spatial scales. (A) Net wind flow across the study area during the flowering season (June–Oct 2015–2022), from HRRR wind modeled with windscape. Shading shows the net flow rate and arrows its direction, with length proportional to rate; the black line is the coastline. Circles are population centroids, colored as in other figures. (B) Pairwise linearized $F_{ST}$ between populations against wind connectivity (log mean least-cost wind flow between each pair, with travel over water down-weighted by 90%). Fill and outline colors identify the two populations in each pair. Open circles are pairs involving Dangermond, whose $F_{ST}$ values are inflated by low sequencing coverage; their outline identifies the other population. (C) Net wind flow within Tranquillon Mountain, downscaled with WindNinja at 100 m with spatially varying vegetation, diurnal slope flows and non-neutral atmospheric stability. Shading and arrows as in A; white circles are individual plants. Across occupied cells, net flow ranges from 1.6 to 3.6 m s$^{-1}$ and its direction spans 44°, compared with 2.0–4.7 m s$^{-1}$ and 52° across the six population sites. (D) Pairwise linearized $F_{ST}$ between Tranquillon Mountain individuals ($n$ = 276) against wind connectivity, with a least-squares fit line. Because wind connectivity declines with distance, panels B and D include isolation by distance; statistical tests control for geographic distance.

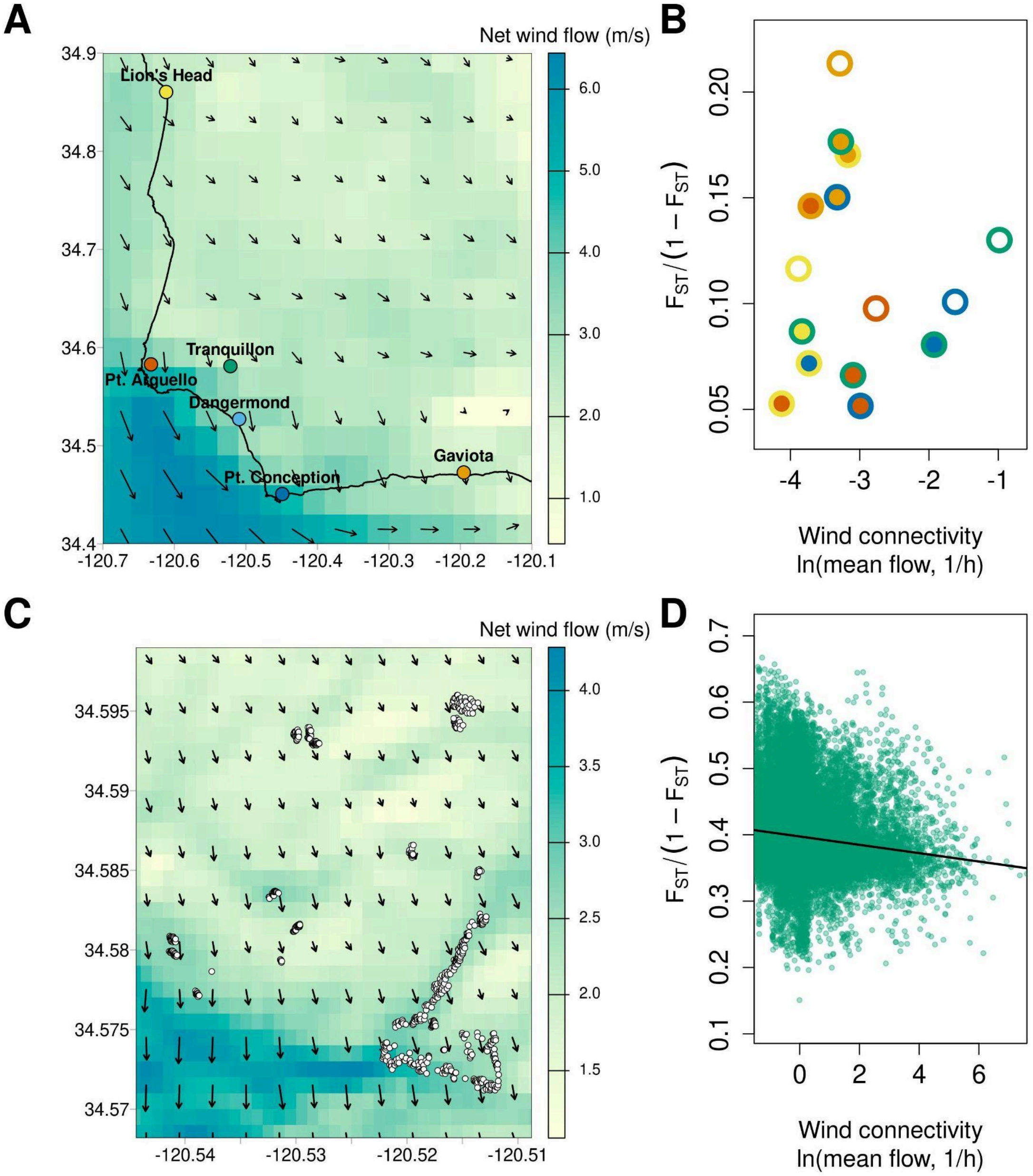

**Figure S7.** Manhattan plots comparing outlier peaks and annotated genes between (A) fog, (B) summer precipitation, and (C) precipitation seasonality. Notice that most peaks are shared across two or more of these hydric variables.

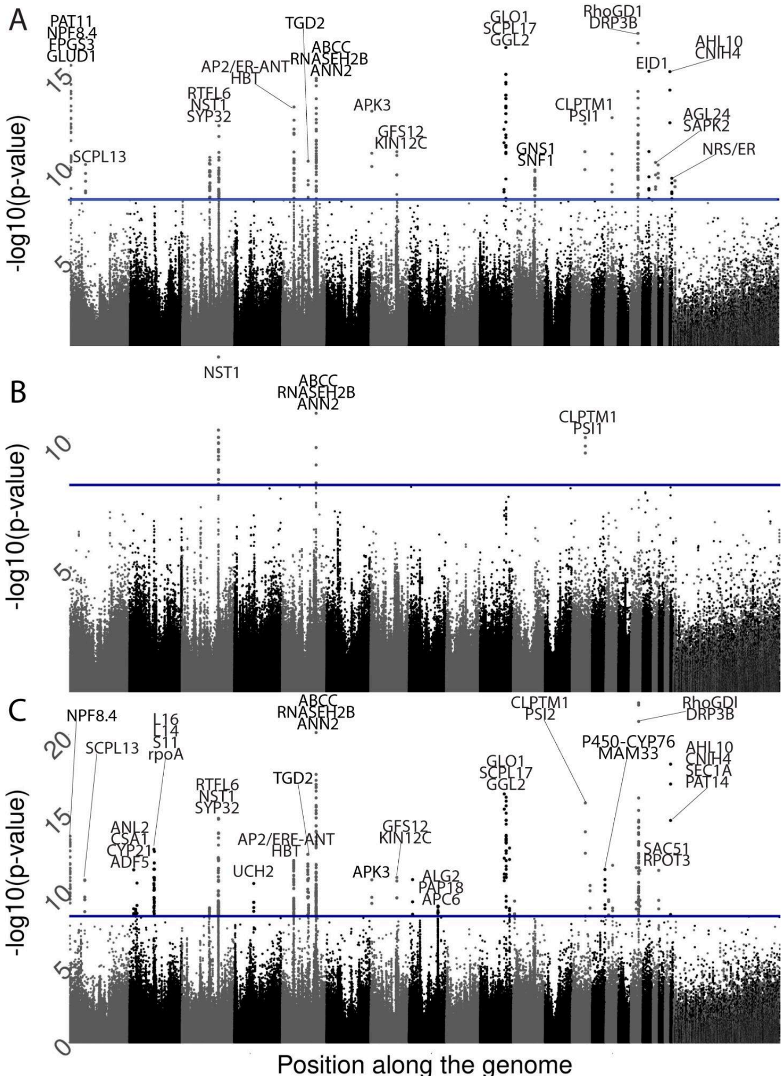


**Figure S8.** Candidate gene overlap across fog and low lying cloud cover (fog), precipitation seasonality (BIO15), and summer precipitation (BIO18). This overlap exceeds the expectation under independent sampling from the genome-wide gene complement (*n* = 33,257 annotated genes): 12 shared genes between precipitation seasonality and summer precipitation against an expectation of 0.05 (hypergeometric test, $p = 5.2 \times 10^{-28}$), 62 shared between precipitation seasonality and fog against an expectation of 0.30 ($p = 4.3 \times 10^{-141}$), and 11 shared between summer precipitation and fog against an expectation of 0.04 ($p = 3.1 \times 10^{-26}$).

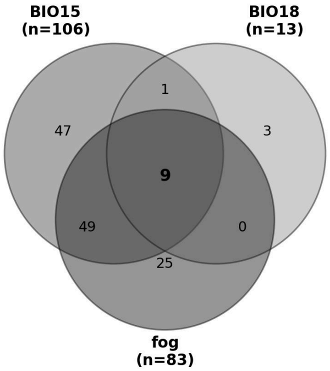

**Table S1.** Outlier genes relative to fog exposure, precipitation seasonality (BIO15), and summer precipitation (BIO18), to accompany the Manhattan plot above (Fig. S9). Since many variants are shared across multiple environmental variables, the first column indicates the variables associated with each gene using a colored circle: light blue for fog, dark blue for summer precipitation, and orange for precipitation seasonality. Variant types are coded as follows: MS = missense variant, SYN = synonymous variant, IG = intergenic variant; DWN = downstream gene; UP = upstream gene; INT = intronic variant; intragenic = IA; and UTR = five prime UTR.

| Variable | Acronym | Position | Min. P-val | Variant Types | My functional/adaptive description |
|---|---|---|---|---|---|
| fog | PAT11 | DEINVI_7: 104622662-104630080 | $1.8x10^{-10}$ | IG | A vacuolar-membrane-localized palmitoylation enzyme that helps plants process xenobiotics. |
| fog, BIO15 | NPF8.4 | DEINVI_7: 111606559-111611204 | $1.7x10^{-13}$ | MS; SYN | A tonoplast-localized glycerate transporter that links photorespiration to nitrogen availability & carbon-nitrogen balance & plays a role in stress response. |
| fog | FPGS3 | DEINVI_7: 114291568-114300508 | $5.2x10^{-9}$ | DWN; IG | Catalyzes conversion of folates to polyglutamate derivatives--important substrates for enzymes that are involved in purine, pyrimidine & amino acid synthesis. Expression increases in response to heat, drought, DNA damage, & oxidative stress. |
| fog | GLUD1 | DEINVI_7: 114303720-114313907 | $5.2x10^{-9}$ | IG; UP | Codes for a major seed storage protein whose overall abundance/composition shifts under drought stress. |
| fog, BIO15 | SCPL13 | DEINVI_7: 34420805-34433981 | $8.7x10^{-11}$ | INT | Produces fibers in plants, & determines traits like cell elongation & locule count. Important in regulating vegetative growth, including of roots. Expression is impacted by environmental stress & the intensity of UV-B radiation. |
| fog, BIO15 | ANL2 | DEINVI_27: 9632399-9637076 | $9.3x10^{-10}$ | DWN; IG | Impacts accumulation of cutins--waxy polymers that are the main component of the plant cuticle. Regulates cell wall mechanical properties. Plays a role in root development (particularly root hairs) & anthrocyanin accumulation. Preferentially expressed in epidermal or subepidermal cells. |
| BIO15 | CSA1 | DEINVI_27: 14017414-14019130 | $2.9x10^{-9}$ | DWN; IG | A plant disease-resistance protein, participating in crosstalk between pathogen-defense & abiotic-stress signaling. |
| fog, BIO15 | CYP21 | DEINVI_27: 16845552-16850193 | $2.3x10^{-9}$ | DWN; IG | Golgi-resident peptidyl-prolyl isomerases that regulate ABA-signaling gene expression during seed germination/seedling growth. Expression impacts oxidative & salinity stress tolerance. |
| fog, BIO15 | ADF5 | DEINVI_27: 16835870-16838755 | $2.3x10^{-9}$ | IG; UP | Promote stomatal closure though the regulation of actin cytoskeleton remodeling. Demonstrated impacts on water loss & plant survival through drought stress. |
| BIO15 | L16 | DEINVI_27: 54145094-54145453 | $2.9x10^{-13}$ | DWN; IG | Ribosomal protein, known to be upregulated or undergo protein modification under drought. |
| BIO15 | L14 | DEINVI_27: 54144617-54144985 | $2.9x10^{-13}$ | DWN | Ribosomal protein that impacts the drought & stress responsiveness of seedlings, purportedly through the induction of antioxidant enzymes. Strong expression difference under salt stress. |
| BIO15 | S11 | DEINVI_27: 54142929-54143339 | $2.9x10^{-13}$ | DWN | Chloroplast translation machinery is broadly affected by drought -- chloroplast ribosomal protein transcripts drop sharply in drought-stressed leaves in other species. |
| BIO15 | rpoA | DEINVI_27: 54141841-54142848 | $2.9x10^{-13}$ | DWN; IG | Chloroplast translation/transcription machinery that catalyzes the transcription of DNA into RNA. PEP-subunit transcripts respond to ABA, & chloroplast ribosomal protein transcripts drop sharply in drought-stressed leaves. |
| fog, BIO15 | RTFL6 | DEINVI_10: 80694686-80708276 | $2.4x10^{-10}$ | IG | Small peptides that negatively regulate cell proliferation, affecting leaf & root length. Involved in shade avoidance syndrome, induced by low red to far right red light ratios. |
| fog, BIO15, BIO18 | NST1 | DEINVI_10: 80925879-80929458 | $2.8x10^{-12}$ | DWN; IG; SYN; UP | The master lignin transcription factor. Regulates secondary cell wall thickening in woody tissues. Is hypothesized to contribute to a desiccation-tolerance strategy by allowing plants to differentiate water-conducting & reinforced, woody supporting tissues. Responds to stress-response signalling. |

| | | | | | |
|---|---|---|---|---|---|
| fog, BIO15 | SYP32 | DEINVI_10: 80569762-80577008 | $1.9x10^{-10}$ | IG; INT; MS; UP | Plays a role in pollen development by modulating protein trafficking & Golgi structure. |
| BIO15 | UCH2 | DEINVI_25: 43217787-43220949 | $4.7x10^{-11}$ | IG; MS; UP | Involved in the release of ubiquitin attached via both peptide & isopeptide linkages, enforcing maintenance of the period of the circadian clock & affecting shoot architecture |
| fog, BIO15 | AP2/ ERF-ANT | DEINVI_17: 26057859-26062490 | $3.7x10^{-13}$ | DWN; IG; INT; UP | Regulates ovule development & floral organ growth. |
| fog, BIO15 | HBT | DEINVI_17: 25620239-25634345 | $1.9x10^{-11}$ | IG | Component of the anaphase promoting complex/cyclosome (APC/C), which controls progression through mitosis & the cell cycle. Essential for meristem differentiation, & particularly the development of the root meristem. |
| fog | TGD2 | DEINVI_17: 56439469-56448202 | $3.7x10^{-13}$ | IG | Component of a phosphatidic acid/lipid transport complex in the chloroplast envelope. Involved in lipid transfer from the endoplasmic reticulum (ER) to plastids. Necessary for thylakoid membrane formation-- the membrane in the chloroplast that hosts photosynthesis. |
| BIO15 | ABCC | DEINVI_17: 70271336-70294689 | $1.0x10^{-9}$ | IG | Localized to the tonoplast, which separates the inner cell sap from the outer cytosol. Regulates the movement of ions & molecules to maintain turgor pressure & regulate stomatal aperture. Mediates vacuolar sequestration of ABA-glucosyl ester & other stress-related metabolites. Expression induced by drought. |
| BIO15 | RNASEH2B | DEINVI_17: 74632367-74638708 | $8.0x10^{-11}$ | IG | Participates in DNA replication. Mediates the excision of single ribonucleotides from DNA:RNA duplexes. Removes misincorporated ribonucleotides from DNA to maintain genome integrity. |
| fog, BIO15 | ANN2 | DEINVI_17: 76779386-76781640 | $4.2x10^{-9}$ | IG | Required for post-phloem sugar transport in root tips to affect primary root growth. Mediates Ca2+ & ROS signaling in stress responses, including stomatal & osmotic responses. |
| fog, BIO15 | APK3 | DEINVI_3: 2810816-2815004 | $1.2x10^{-13}$ | IG; UP | Assists plants with the catalysis of sulfate (which is deposited when sulfate-rich sea spray nucleates coastal clouds/fog), & is essential for plant viability & reproduction. |
| fog, BIO15 | GFS12 | DEINVI_3: 58413617-58423042 | $1.8x10^{-11}$ | IG; UP | Transports proteins to seed storage vacuoles. |
| fog, BIO15 | KIN12C | DEINVI_3 :57961242-57972045 | $1.8x10^{-11}$ | IG | Involved in the spatial control of cytokinesis. Kinesin-family microtubule motors show drought/water-deficit-responsive expression in multiple species, consistent with microtubule dynamics supporting stress-related structural remodeling. |
| BIO15 | APC6 | DEINVI_12: 63521968-63528459 | $6.7x10^{-11}$ | IG | Component of the anaphase promoting complex/cyclosome (APC/C), which controls progression through mitosis & the cell cycle. Influences leaf size, vein patterning, vascular bundle size, lateral root number, & root elongation. |
| BIO15 | ALG2 | DEINVI_12: 63763759-63767234 | $6.7x10^{-11}$ | IG; INT; UP | Adds mannose residues during lipid-linked oligosaccharide biosynthesis (N-glycosylation pathway). A related plant glycosylation enzyme (ALG10) is required for normal leaf development. |
| BIO15 | PAP18 | DEINVI_12: 64070841-64074959 | $1.4x10^{-9}$ | IG | Responsive to drought & phosphate-deficiency stress. Mobilizes phosphate to support growth under adverse conditions. |
| fog, BIO15 | GLO1 | DEINVI_8: 56546210-56549123 | $2.8x10^{-10}$ | DWN; IG | A committed enzyme of photorespiration. Involved in a pathway documented to support plant resistance to abiotic stresses including drought & high light by managing excess energy & H2O2 signaling. |
| fog, BIO15 | SCPL17 | DEINVI_8: 62871050-62887538 | $1.0x10^{-10}$ | INT | Involved in the biosynthesis of glucosinolates (GSLs), nitrogen- & sulfur-containing secondary metabolites derived from amino acids that protect plants from pests & in some plants are involved in auxin biosynthesis |
| fog, BIO15 | GGL2 | DEINVI_8: 65124349-65129807 | $2.3x10^{-13}$ | DWN; IG; INT | Play a role in stomatal dynamics, density, & morphology. |
| fog, BIO15 | LIP1 | DEINVI_6: 48254126-48262478 | $1.8x10^{-12}$ | IG; INT | A key regulator of the circadian clock & photomorphogenesis, influenced by the spectrum of light frequencies. |
| fog | GNS1 | DEINVI_6: 49099793-49104495 | $1.8x10^{-9}$ | IG; UP | A pathogenesis-related enzyme that hydrolyzes the beta-1,3-glucan of fungal cell walls. Induced upon pathogen challenge. |
| fog | SNF1 | DEINVI_6: | $4.5x10^{-10}$ | DWN; IG; | Subunit of the SnRK1 energy-sensing kinase complex. Activated by low-energy/stress conditions. Downregulates |

| | | | | | |
|---|---|---|---|---|---|
| | | 49121438-49124049 | | INT | ATP-consuming biosynthesis, while upregulating catabolic energy-generating processes. SnRK1 expression enhances drought tolerance. |
| fog, BIO15, BIO18 | CLPTM1 | DEINVI_19: 29765991-29771035 | $6.0x10^{-13}$ | DWN; IG | In *Arabidopsis* lines with mutant trichomes, this transmembrane protein is among the top 25 genes with strong differential expression. Downregulated in response to water stress & heat. |
| fog, BIO15 | PSI2 | DEINVI_19: 30052184-30057752 | $3.0x10^{-12}$ | UTR; IA | Promotes seedling & growth probably via the regulation of peptide phytohormones acting as growth factors. Highly sensitive to the spectrum & intensity of light frequencies |
| fog, BIO15 | MAM33 | DEINVI_14: 28675172-28677667 | $3.2x10^{-12}$ | IA; INT | In yeast, MAM33 ensures the proper assembly of ribosomes, but its role in plants is not yet clear. |
| fog, BIO15 | P450-CYP76 | DEINVI_14: 28681828-28683863 | $3.2x10^{-12}$ | UP | Metabolizes with high efficiency a wide range of xenobiotics. Expression strongly impacted by drought & salt stresses. |
| fog, BIO15 | RhoGDI | DEINVI_20: 16884617-16888352 | $2.3\times10^{-14}$ | INT; STN; MS | Negative regulators of RHO GTPases, which in turn control various cellular processes, from cell adhesion to migration to proliferation |
| fog, BIO15 | DRP3B | DEINVI_20: 17300485-17309354 | $8.5x10^{-14}$ | DWN; IG | Involved in the control of mitochondrial & peroxisomal division & morphology. |
| fog, BIO15 | EID1 | DEINVI_15: 15365141-15365905 | $1.7x10^{-9}$ | DWN; IG | Negatively regulates phytochrome A (phyA)-specific light signaling.Targets activated phyA-pathway components for ubiquitin-dependent degradation & shifting red/far-red light responsiveness, modifying the phase & period of a plant's circadian rythm, & affecting processes like morphogenesis & flowering time. |
| fog | AGL24 | DEINVI_2: 13318019-13340714 | $9.1x10^{-11}$ | INT | Affects the timing of vegetative-to-reproductive floral transition in response to photoperiod, promotes inflorescence identity, & regulates root growth. |
| BIO15 | SAC51 | DEINVI_2: 13220955-13225308 | $6.6x10^{-12}$ | DWN | A bHLH transcription factor. Regulates xylem differentiation, impacting stem & root elongation. |
| BIO15 | RPOT3 | DEINVI_2: 13211416-13218760 | $6.6x10^{-12}$ | IA; MS | A phage-type nucleus-encoded polymerase (NEP) enzyme. Required for chloroplast development & leaf mesophyll cell proliferation. |
| fog | SAPK2 | DEINVI_2: 13068550-13075163 | $2.3x10^{-9}$ | INT | Plays an important role in ABA-mediated seed dormancy. Reduces water loss by driving the accumulation of compatible solutes, closing stoma, & increasing the expression of stress-response & slow anion channel-associated genes. Promotes ROS-scavenging. Confers salt, drought, & other hyperosmotic stress tolerance. Expressed particularly in the leaves & roots. |
| fog, BIO15, BIO18 | AHL10 | DEINVI_21: 349838-351884 | $9.8x10^{-16}$ | IG; UP | Transcription factor that specifically binds AT-rich DNA sequences related to the nuclear matrix attachment regions (MARs). Regulates growth, particlarly of roots. Significantly upregulated under drought & salt stress, enhancing drought tolerance. |
| fog, BIO15, BIO18 | CNIH4 | DEINVI_21: 345599-348147 | $9.8x10^{-16}$ | DWN | Cornichon proteins are COPII cargo receptors that escort transporters from the ER to the Golgi/plasma membrane. Best characterized in pollen tube Ca2+ channel trafficking & auxin transporter localization. |
| BIO15 | SEC1A | DEINVI_21: 331989-340165 | $4.9x10^{-9}$ | INT | Promote pollen tube growth & pollen germination. |
| BIO15 | PAT14 | DEINVI_21: 324867-329700 | $4.9x10^{-9}$ | UP | A Golgi protein that plays a role in leaf senescence & immunity gene expression via palmitoylation. |
| fog | NRS/ER | DEINVI_21: 4566259-4570150 | $4.7x10^{-10}$ | DWN; IG | Helps build rhamnose, a core building block of cell walls, & produce protective plant compounds like flavonoids. |

**Figure S9.** Enriched gene ontology terms associated with (A) fog & low-lying cloud cover, (B) precipitation seasonality & (C) summer precipitation, for biological processes (BP), molecular function (MF), & cellular components (CC). There were three identical terms shared between multiple variables: cellulose synthase activity, diterpene phytoalexin biosynthetic process, and protein localization to the cell surface. Two additional term pairs were near-misses— closely related but not identical GO terms describing the same underlying process: wax biosynthesis regulation (positive regulation of wax biosynthetic process in fog; regulation of wax biosynthetic process in warm-season precipitation) and polysaccharide catabolism (polysaccharide catabolic process in fog; glucan catabolic process in precipitation seasonality).

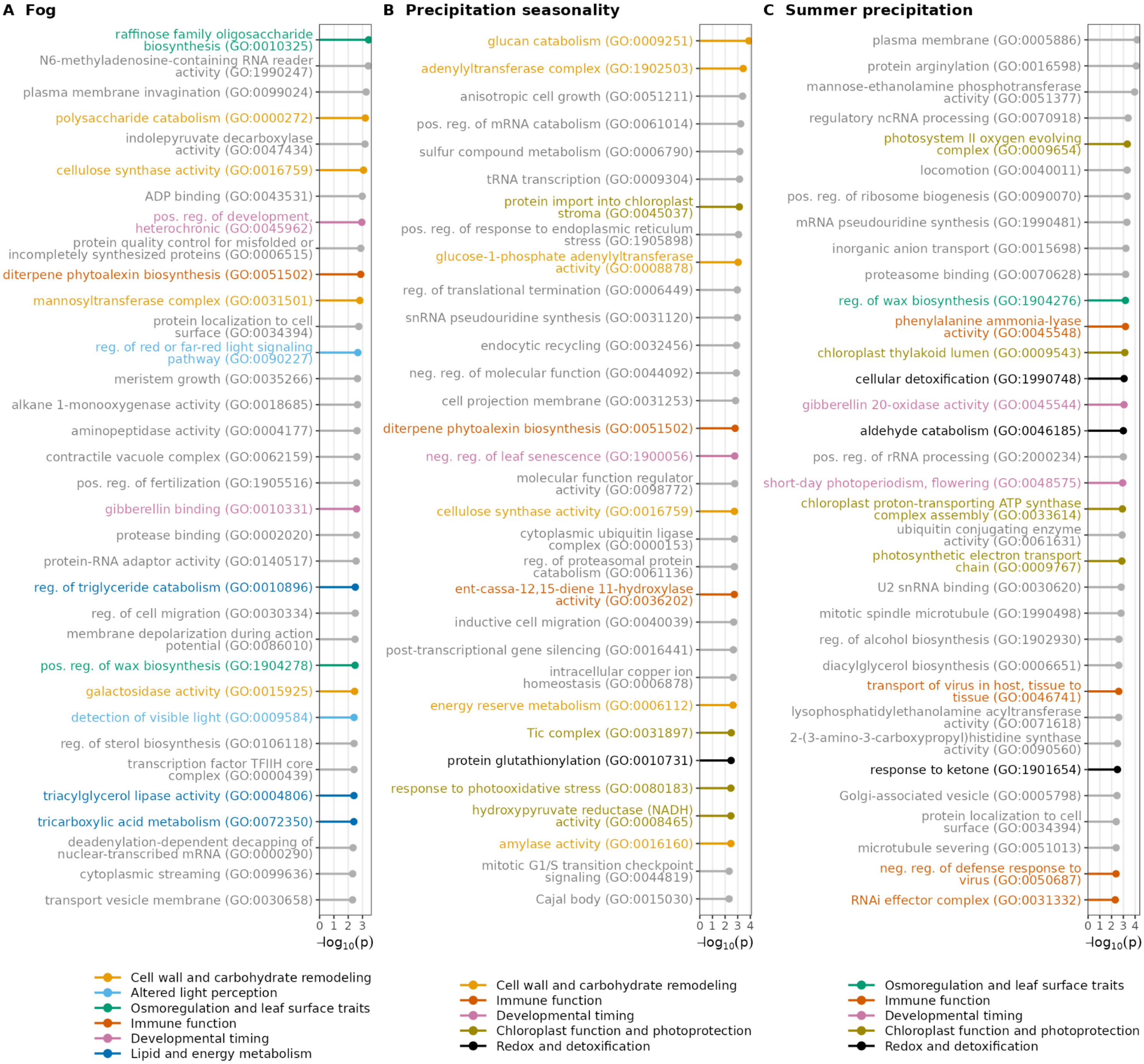

**Figure S10.** Tissue expression of candidate adaptation loci. Transcripts per million (TPM) for 51 candidate genes from GxE on fog, summer precipitation, and precipitation seasonality. TPM were computed across eight RNA-seq libraries from one population, spanning root (n=4), shoot (n=3), and leaf (n=1) tissue from four individuals (1, 2, 5, and M). Cell color reflects relative tissue pattern within a gene and is not comparable in magnitude across genes: it shows each gene's own log2(TPM+1) value min–max scaled across its eight samples (row-normalized). Note that because leaf tissue is represented by a single individual and no formal differential-expression test was applied, tissue differences shown here are descriptive rather than statistically supported.

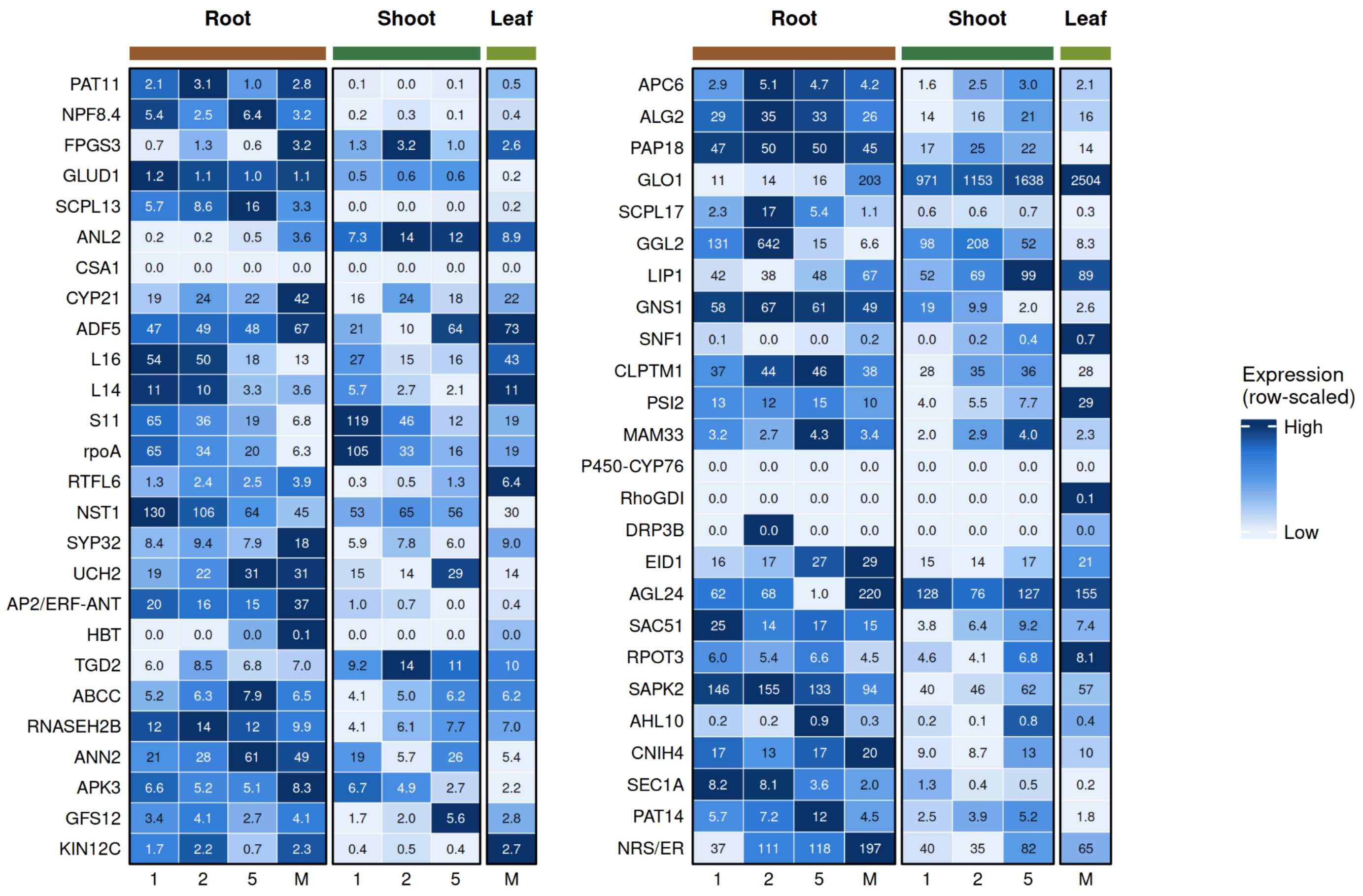

**Figure S11**. Fog-associated outlier peaks for each of 12 major scaffolds and their position relative to inferred centromeres (grey box). GxE fog p-values are shown first, followed by gene density and LINE enrichment. Dashed lines in the lower two tracks give genome-wide means. At the bottom of each panel are syntenic gene pairs between DEINVI_00006 and lettuce (*Lactuca sativa*). x is the gene's position on DEINVI_00006 and y is its partner's position on the partner chromosome (only chromosomes with ≥ 20 pairs shown), such that collinear blocks are shown as diagonal lines. DEINVI_00006 is the only chromosome-scale scaffold whose centromere cleanly separates segments of two different lettuce chromosomes, though on three others a single gene of the left-arm partner lies just past the centromere before the partner changes, suggesting a nearby junction. The fog peak lies ~350 kb from the first syntenic gene of the right ancestral segment.

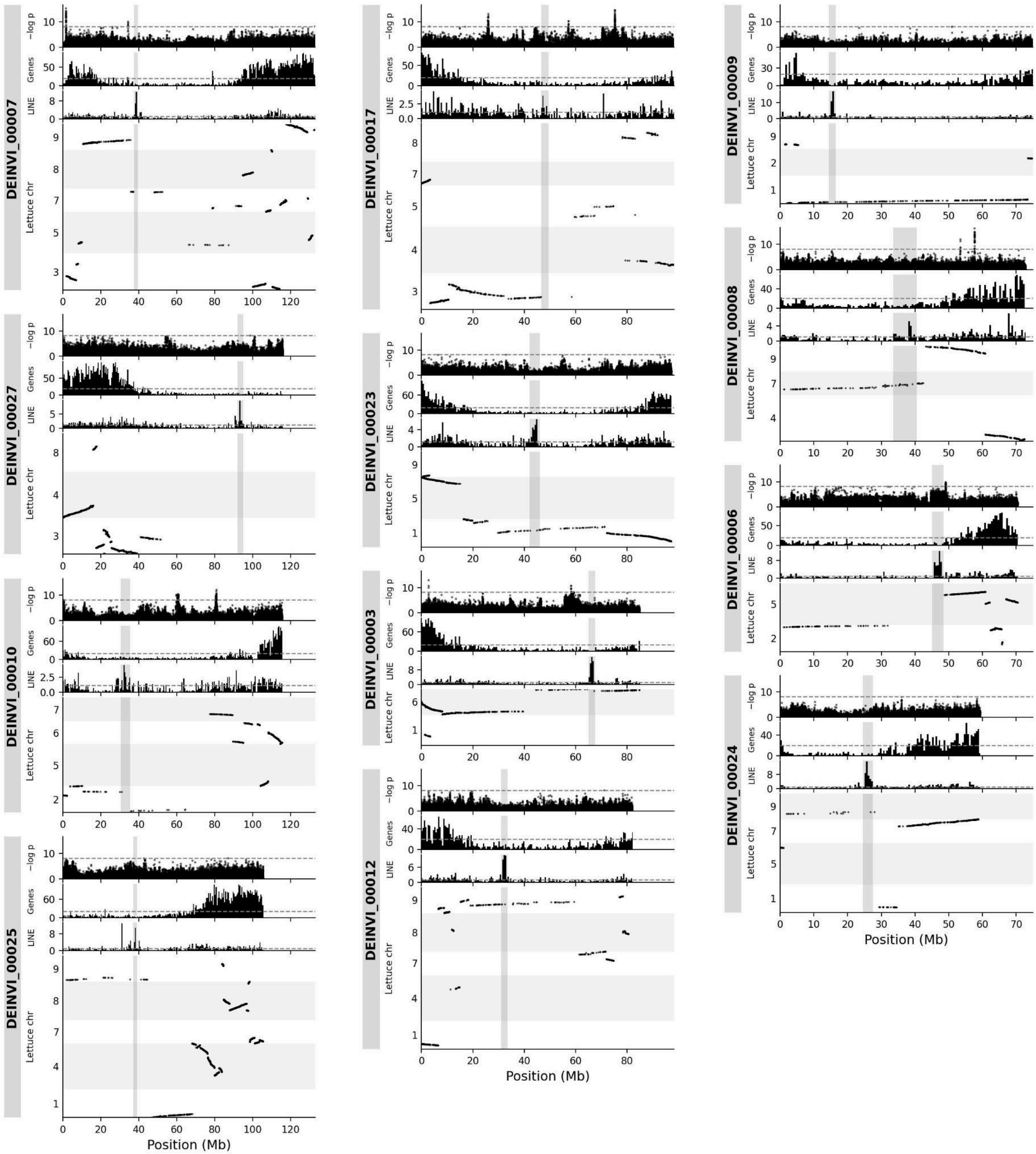

**Figure S12**. Population-genomic outliers, structure, & synteny along chromosome DEINVI_00006. The grey box marks the putative centromere. (A) GxE association with long-term fog. The dashed line is the Bonferroni threshold. (B) Sliding-window PC1, one line per individual. Lines are colored by individuals' long-term fog (days/month). At most 15 randomly chosen individuals are drawn per locality. Note that the centromere carries two non-recombining haplotypes (A & B, with 649 diagnostic SNPs). Relating this to **Fig. 3B**, LD is moderate between the fog outlier haploblock and centromeric haploblock because the favourable fog haplotype occurs 33% of the time on A and 15% of the time on B. (C) Proportion of heterozygous sites per window, one line per individual. (D, E) Syntenic gene pairs between DEINVI_00006 and sunflower (*Helianthus annuus*) and lettuce (*Lactuca sativa*). x is the gene's position on DEINVI_00006 and y is its partner's position on the partner chromosome (only chromosomes with ≥ 20 pairs shown), such that collinear blocks are shown as diagonal lines.

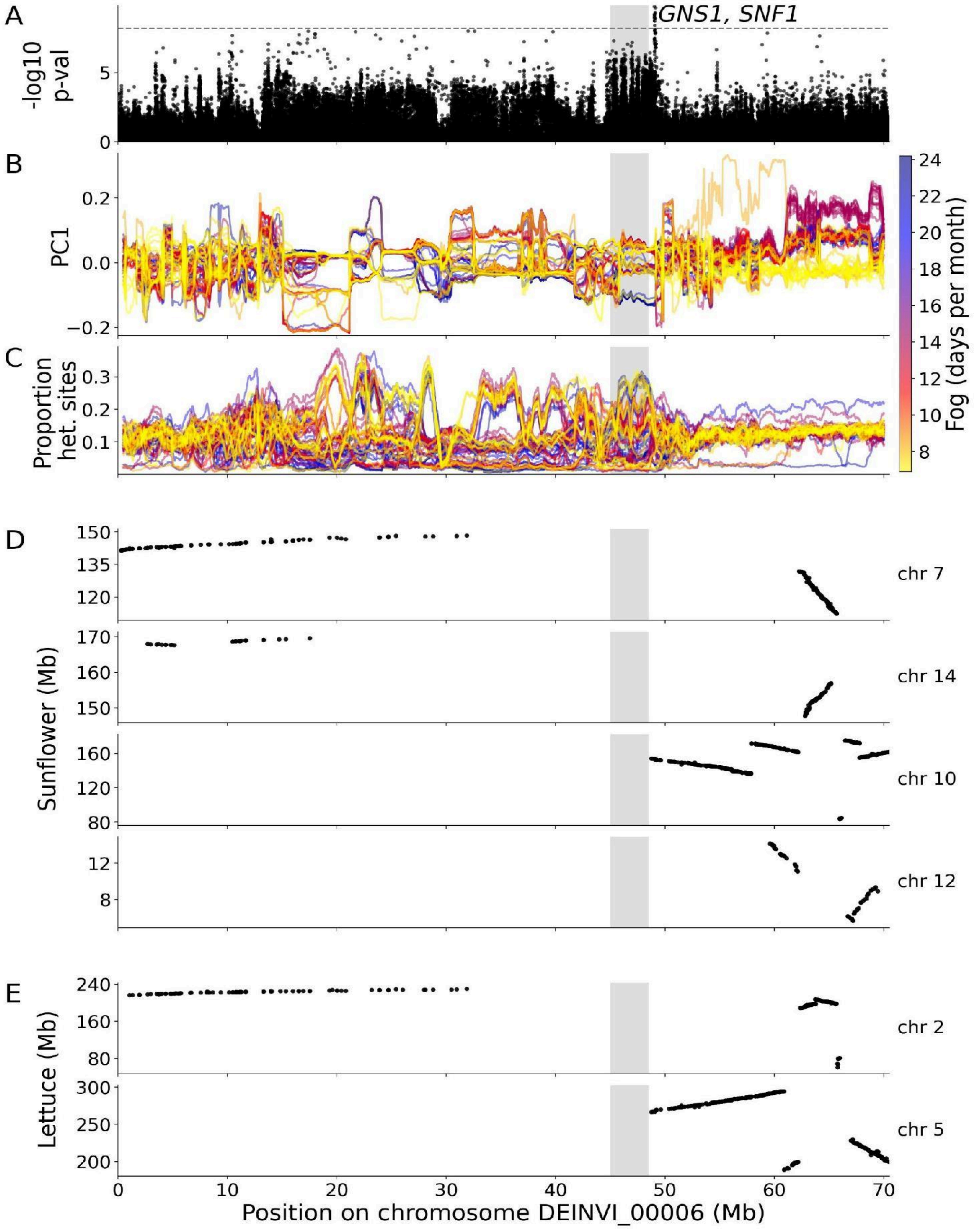

**Figure S13**. Frequency of the centromeric-associated haplotype in chromosome DEINVI_00006 between germination years.

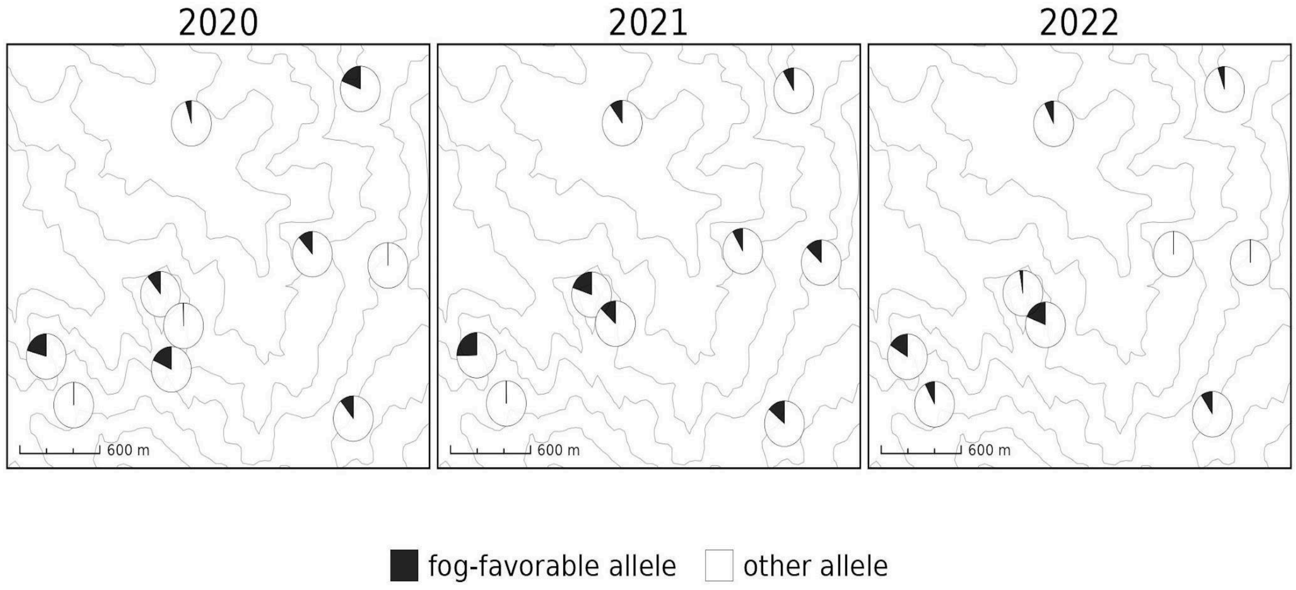


**Figure S14.** Tajima's D at the DEINVI_00006 fog peak and other fog outlier peaks, compared with random genomic blocks. Tajima's D was estimated per population in sliding 100-SNP windows. Every value shown is the median Tajima's D of the windows overlapping a region. Boxes show the median and interquartile range; whiskers show 1.5 × IQR. On the right, the grey boxes show 3,000 random blocks at the same distance from their own centromere as the DEINVI_00006 peak, and the diamonds show the DEINVI_00006 peak. On the left, the white boxes show 3,000 random 115-kb blocks placed anywhere on the chromosome-scale scaffolds outside of outlier peaks, and points show the other 30 outlier peaks.

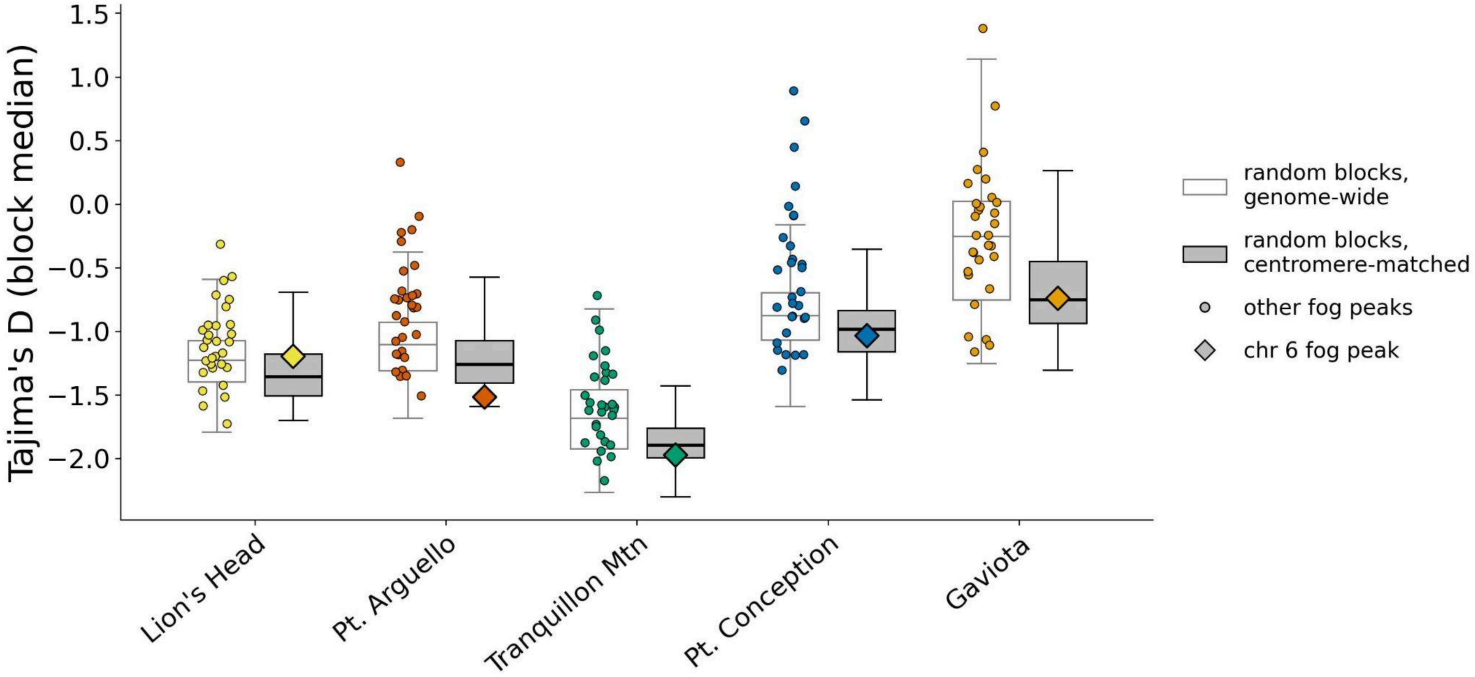

**Supplementary Note. The centromeric histone CENH3.** Under the centromere-drive model, centromeres that gain a transmission advantage in asymmetric female meiosis spread at the expense of organismal fitness, and centromeric proteins such as CENH3 (the centromere-specific histone H3 variant) evolve adaptively to suppress them. Recurrent, rapid selection in CENH3 is therefore an expected footprint of suppressor evolution, for example in *Mimulus* being concentrated in the N-terminal tail (Finseth *et al.* 2015, Zedek & Bureš 2016), so we examined CENH3 for that footprint. Indeed, *D. increscens* has a single CENH3 gene in the assembly, on DEINVI_00019, whose histone-fold domain is strongly conserved but whose N-terminal tail evolves unusually fast. tblastn searches with the tail found no second copy, suggesting that the two CENH3 paralogues of *H. annuus* and *H. anomalus* arose after the Helianthus lineage split from *D. increscens*. Across the whole protein, *D. increscens-H. annuus* dN/dS (0.065-0.084) is lower than for 85-91% of 1,995 one-to-one orthologues, reflecting the histone fold (dN/dS 0.07-0.13). In contrast, the tail's dN/dS against five Helianthus sequences (0.39-0.57) exceeds that of 94-99% of genes, and between *D. increscens* and *Layia glandulosa*, the other Madiinae examined, it is 1.36 (versus 0.22 for the histone fold). The tail is also longer in both Madiinae, through a shared expansion of a repeated motif. Codon-model tests on nine sequences detected no positively selected sites or positive selection on the *D. increscens* branch, but have little power with so few sequences, and relaxed constraint on the tail cannot be excluded.

**Table S2.** Top three models, by AIC, resulting from model comparison of how fog, summer precipitation (BIO18), and their interaction might contribute to the proportion of ray florets and the height to width ratio.

| Phenotype | N | Formula | Model weight | AIC | Delta AIC | Variable | p-value | Estimate ± std error | z/t-score |
|---|---|---|---|---|---|---|---|---|---|
| Proportion ray florets | 138 | Fog + BIO18 | 0.25 | -387.82 | 0 | intercept<br>Fog<br>BIO18 | $7.60 \times 10^{-112}$<br>0.04<br>0.04 | -0.47 ± 0.02<br>0.05 ± 0.03<br>0.07 ± 0.03 | -22.47<br>2.10<br>2.05 |
| | | Fog + BIO18 + CWD | 0.165 | -386.98 | 0.83 | (Intercept)<br>Fog<br>BIO18<br>CWD | $3.80 \times 10^{-113}$<br>0.03<br>0.02<br>0.25 | -0.47 ± 0.02<br>0.05 ± 0.03<br>0.08 ± 0.04<br>0.02 ± 0.02 | -22.61<br>2.19<br>2.33<br>1.16 |
| | | Fog * CWD | 0.148 | -386.76 | 1.05 | (Intercept)<br>Fog<br>CWD<br>Fog x CWD | $2.50 \times 10^{-99}$<br>0.3<br>0.54<br>0.02 | -0.49 ± 0.02<br>0.02 ± 0.02<br>0.01 ± 0.02<br>0.10 ± 0.04 | -21.15<br>1.03<br>0.62<br>2.29 |
| Plant height:width | 115 | Fog + BIO18 | 0.42 | 307.01 | 0 | intercept<br>Fog<br>BIO18 | 0.24<br>$1.40 \times 10^{-4}$<br>0.02 | 0.90 ± 0.76<br>-0.67 ± 0.18<br>-2.04 ± 0.89 | 1.18<br>-3.80<br>-2.30 |
| | | Fog + BIO18 + CWD | 0.138 | 309.23 | 2.23 | Intercept<br>Fog<br>BIO18<br>CWD | 0.25<br>0.00033<br>0.024<br>0.96 | 0.89 ± 0.768<br>-0.67 ± 0.19<br>-2.04 ± 0.90<br>0.004 ± 0.09 | 1.16<br>-3.59<br>-2.26<br>0.05 |
| | | Fog * BIO18 | 0.14 | 309.23 | 2.23 | intercept<br>Fog<br>BIO18<br>Fog x BIO18 | 0.24<br>0.03<br>0.02<br>0.97 | 0.90 ± 0.76<br>-0.66 ± 0.30<br>-2.03 ± 0.89<br>0.02 ± 0.36 | 1.18<br>-2.20<br>-2.28<br>0.04 |

**Figure S15.** Phylogenetic and geographic context for the Gaviota tarplant (*Deinandra increscens* subsp. *villosa*). (A) The closest relative of Gaviota tarplant is the Grassland tarweed (*Deinandra increscens* subsp. *villosa*). (B). Gaviota tarplant has a fragmented and very coastal distribution in Central California.

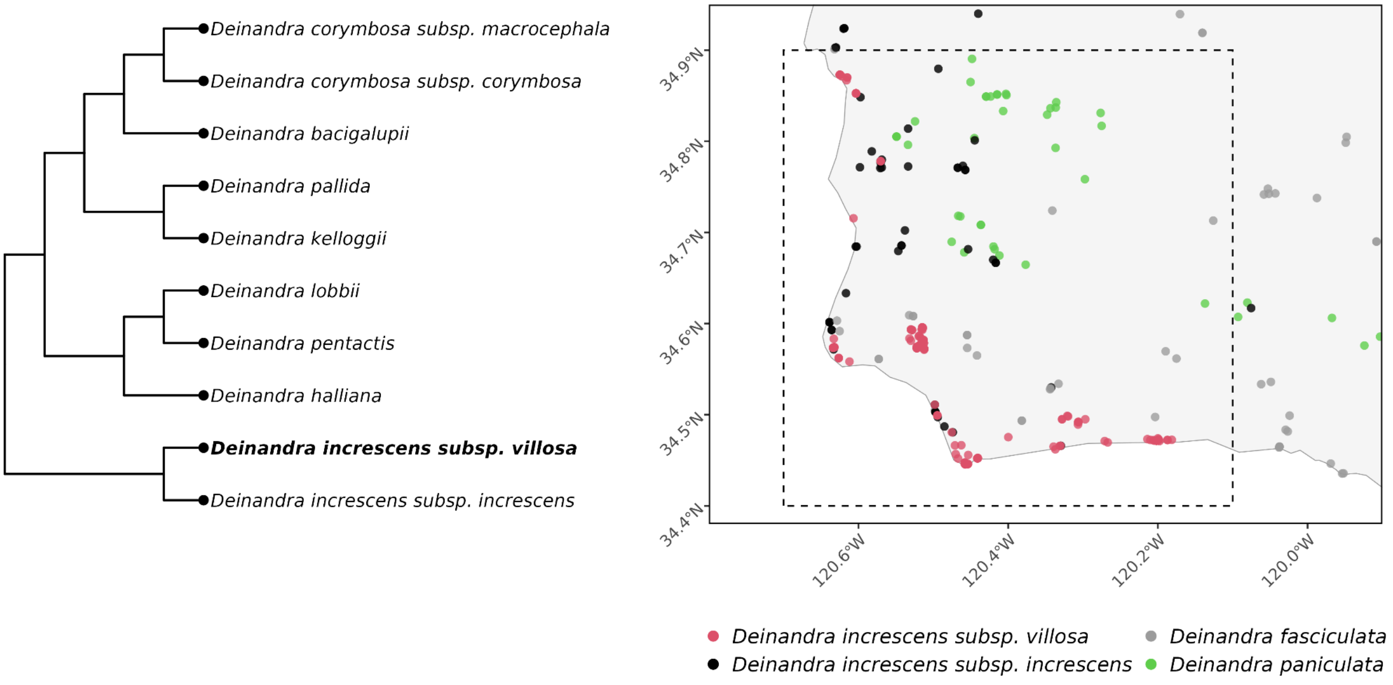


**Figure S16.** Collinearity and PCA of retained environmental variables for genotype by association analyses. The left plot shows Spearman's correlation coefficients between retained variables. The right plot is the PCA, showing how these retained variables split up the ecological space occupied by tarplant populations, with population colors assigned as elsewhere: Lion's Head in yellow, Pt. Arguello in dark orange; Tranquillon Mountain in green, Gaviota in light orange, Dangermond in light blue, and Pt. Conception in dark blue.

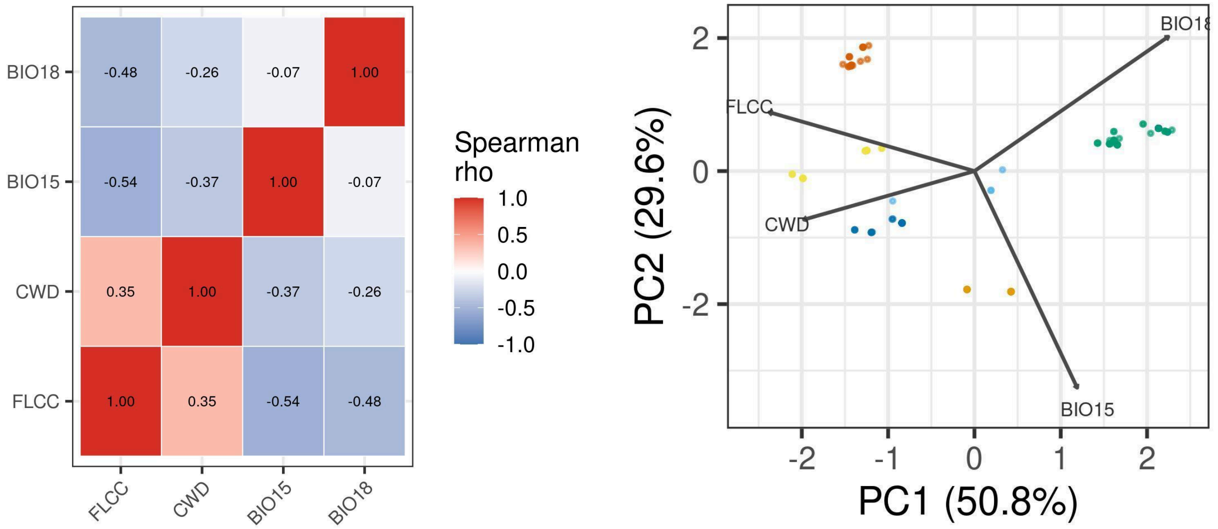

**Figure S17.** Map of new and historic specimens used in phenotype analysis, collected between 1927 and 2024. Colored points indicate individual specimens. Colors represent designated localities.

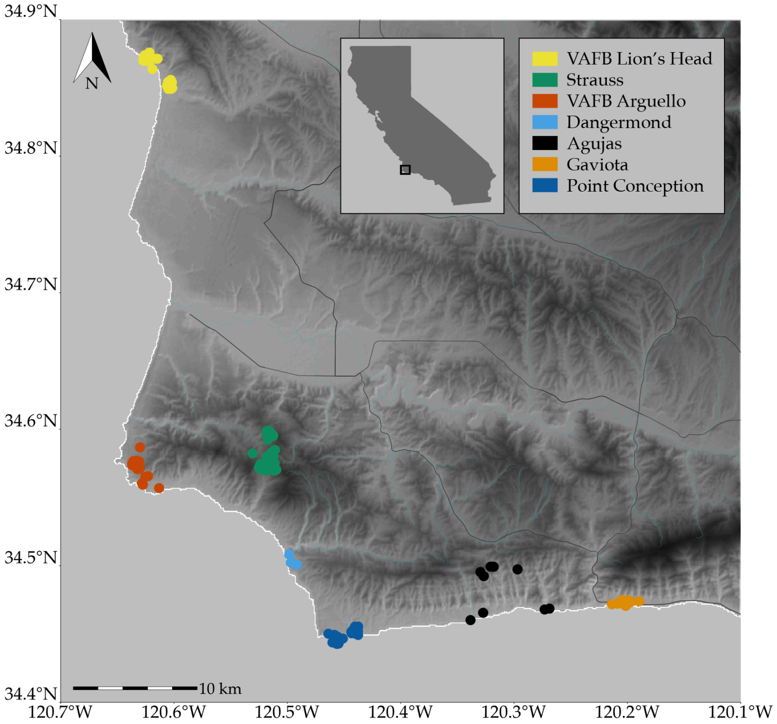


**Figure S18.** Comparison of the two fog metrics used in this publication, pooled over all sampled coordinates and years of overlap between the two products (2000–2014; n = 1,170 coordinate-years). The Werner (MODIS FLCC) product measures mean days per month with fog (June–September); the Kling product measures percent of hours per month with fog over the same season. Despite differing units, the two are strongly correlated (Pearson $r = 0.78$, 95% CI [0.75, 0.80], $t(1168) = 42.3$, $p < 1 \times 10^{-236}$), supporting their joint use as independent cross-validating measures of fog exposure. Line and shaded band show the linear fit and 95% confidence interval.

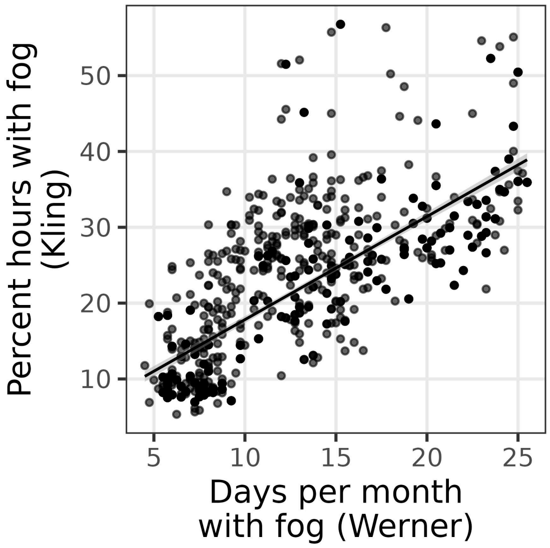
